# A microscopic investigation of the effect of random envelope fluctuations on phoneme-in-noise perception

**DOI:** 10.1101/2022.12.27.522040

**Authors:** Alejandro Osses, Léo Varnet

## Abstract

In this study, we investigated the effect of specific noise realizations on the discrimination of two consonants, /b/ and /d/. For this purpose, we collected data from twelve participants, who listened to the words /aba/ or /ada/ embedded in one of three background noises. All noises had the same long-term spectrum but differed in the amount of random envelope fluctuations. The data were analyzed on a trial-by-trial basis using the reverse-correlation method. The results revealed that it is possible to predict the categorical responses with better-than-chance accuracy purely based on the spectro-temporal distribution of the random envelope fluctuations of the corresponding noises, without taking into account the actual targets or the signal-to-noise ratios used in the trials. The effect of the noise fluctuations explained on average 8.1% of the participants’ responses in white noise, a proportion that increased up to 13.3% for noises with a larger amount of fluctuations. The estimated time-frequency weights revealed that the measured effect originated from confusions between noise fluctuations and relevant acoustic cues from the target words. Substantially similar conclusions were obtained from simulations using an artificial listener. We argue that this token-specific effect of noise is a form of informational masking.

## I. INTRODUCTION

Studies of speech-in-noise perception often rely on speech reception thresholds (SRTs) as a measure of the participant’s intelligibility. An SRT is defined as the signal-to-noise ratio (SNR) at which a specific phoneme, word, or sentence score is achieved for a specific set of speech sounds (e.g., Plomp and Mimpen, 1979). SRTs are typically obtained from many trials to neutralize the response variability that is assumed to be non-relevant for the study (Green, 1964), including the “external variability” introduced by the randomness in the noise stimuli (e.g., Ewert and Dau, 2004). Following the rationale from some existing studies (Jürgens and Brand, 2009; Zaar and Dau, 2015), we will refer to this classical approach as providing a macroscopic view on speech perception performance. The term macroscopic refers to the fact that speech intelligibility is assumed to be based on long-term characteristics of the masking sounds rather than on their detailed waveforms. In contrast, a microscopic view on speech perception is obtained from approaches where intelligibility is assessed on a trial-by-trial basis. For the purposes of this study, specific noise realizations having the same long-term characteristics will be referred to as noise tokens. In this sense, a macroscopic estimate of speech intelligibility will be influenced by an “overall effect” of noise, whereas a microscopic estimate will be related to a “token-specific effect” of noise.

### A. Macroscopic approach: Overall effect of noise

Speech intelligibility measured using macroscopic approaches can be seen as reflecting an overall effect of noise that is not expected to change when different noise tokens are used, as long as enough noise tokens are used during the experiments. As a consequence, this overall effect is related to the fixed, long-term noise statistics and does not depend on the specifics of the noise tokens. Macroscopic intelligibility estimates are usually interpreted in terms of energetic masking (EM) and modulation masking (MM).

The concept of EM refers to a masking effect that occurs when the target and the (undesired) masker sounds overlap in a set of frequency bands (French and Steinberg, 1947). In such cases, the weakest elements in the speech sounds become less audible when the SNR is decreased, causing more and more recognition errors, until the sounds are no longer detected (see, e.g., Li *et al*., 2010, their Fig. 1). The conception of noise as primarily an energetic masker forms the basis for several objective intelligibility metrics, including the articulation index (AI) (French and Steinberg, 1947) and the speech transmission index (STI) (Houtgast and Steeneken, 1985; Payton and Braida, 1999). With these metrics, two noises that have the same long-term distribution of energy with respect to the speech signal are assumed to produce the same EM effect and, hence, the same intelligibility.

**FIG. 1.**
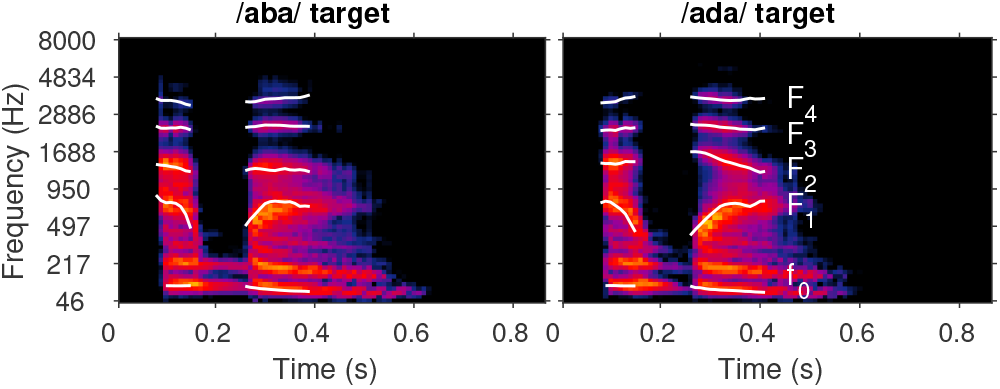
T-F representations of the two targets used in the experiment, /aba/ and /ada/. The time and frequency resolutions are the same as those used for the analysis (see Sec. IIE3a). Lighter regions indicate higher amplitudes in a logarithmic scale. The white traces indicate the fundamental frequency (*f*_0_) and formant trajectories (*F*_1_–*F*_2_).

The concept of MM is similar to EM but operates in the temporal modulation domain. Here, the long-term property that determines the amount of masking is the amplitude of noise envelope fluctuations within each of the analyzed frequency bands. Random envelope fluctuations have been shown to be detrimental to listening performance. As stated by Drullman (1995), they in-duce a “sorting problem” because the weak elements in the signal are likely to be confused with irrelevant fluctuations from the masker. Importantly, this phenomenon is present even for steady-state masking noise, such as white noise or speech-shaped noise, due to the intrinsic random envelope fluctuations present in the masker signal (Dau *et al*., 1999; Stone *et al*., 2011, 2012; Varnet and Lorenzi, 2022). In terms of speech performance,Dubbelboer and Houtgast (2008) and Jørgensen and Dau (2011) proposed that the MM effect is related to the amount of envelope fluctuations in the signal relative to the fluctuations in the noise, in different modulation bands, an idea that was formalized into the signal-to-noise ratio in the envelope domain (SNR_env_).

In summary, the overall impact of noise on intelligibility using the concepts of EM and MM is determined by long-term statistics of the noise masker, based on either band-level energy or the amplitude of random fluctuations, respectively. A macroscopic approach provides an efficient way to measure the impact of EM and/or MM on speech intelligibility, for example by comparing the SRT or the mean intelligibility scores between noise maskers with different long-term characteristics, measured across a large number of noise tokens (e.g., Francart *et al*., 2011; Stone *et al*., 2011).

### B. Microscopic approach: Token-specific effect of noise

A macroscopic approach may not provide a complete picture of the mechanisms involved in speech-in-noise intelligibility, because the averaging operation obscures information about how individual speech tokens are perceived (see, e.g., Singh and Allen, 2012; Zaar and Dau,2015). For instance, the concepts of AI, STI, or SNR_env_ predict that different tokens of the same noise process should be approximately the same. However, this is not the case as it can be shown by adopting a microscopic approach that looks at speech intelligibility at a trial-by-trial level.

More specifically, in speech-in-noise tasks, two noise tokens that have the same long-term statistical properties may not necessarily yield the same intelligibility, as a consequence of the specific sample-by-sample differences between their waveforms. Namely, when a target has to be recognized in a noise background, one particular noise token may mislead the listener while other tokens may not, depending on the specific configuration of random envelope fluctuations in the masker envelope. This phenomenon is referred here to as “token-specific effect” of noise. This microscopic view on speech intelligibility is best illustrated in an experiment by Zaar and Dau (2015), using a fixed set of (frozen) noise tokens. Their participants had to identify consonant-vowel words embedded in white noise at six SNRs. The authors were particularly interested in assessing the effect of various sources of variability in the task. They found that a 100-ms temporal shift of the masking noise waveform could induce a significant perceptual effect, that was well above the assessed within-listener variability. Thus, a given speech utterance was found to be either more or less robust, eliciting a different pattern of confusions, depending on whether the utterance was presented along one specific noise token or its time-shifted version. In their interpretation of this finding,Zaar and Dau noted that “[…] the common assumption in various previous studies of an invariance of consonant perception across steady-state noise realizations cannot be supported by the present study. In fact, the results obtained here suggest that the interaction between a given speech token and the spectro-temporal details of the ‘steady-state’ masking noise waveform matter in the context of microscopic consonant perception. When analyzing responses obtained with individual speech tokens, averaging responses across noise realizations thus appears problematic” (p. 1263).

In order to get a more physiologically-informed insight into the token-specific effect of noise, the variability across tokens may not be considered at the level of the waveform, but rather at the output of the auditory cochlear processing (e.g., Dau et al., 1999). The random envelope fluctuations, that is, the fluctuations in energy induced by the noise at the output of each cochlear channel, vary from one realization to the other, potentially inducing a token-specific effect.

For instance, using steady-noise maskers, Varnet and Lorenzi (2022) showed that the exact temporal distribution of random envelope fluctuations in a trial has a systematic influence on the detection of an amplitude-modulated target. Namely, it can bias the participant’s response towards perceiving a 4-Hz modulation (or not) depending on the token-specific configuration. We found comparable conclusions in two of our previous studies on phoneme discrimination in white noise (Varnet et al., 2013, 2015a). The microscopic approach used in these studies was based on a fine-grained statistical analysis that relates the random envelope fluctuations in a given trial to the corresponding response of the listener. The outcome of this analysis is a time-frequency (T-F) matrix of perceptual weights, named auditory classification image (ACI), which highlights the T-F regions of the stimulus where an increase of random envelope fluctuations induces a systematic bias in the listener’s phonetic decision. These conclusions have not only been obtained for different phonetic contrasts (/b/-/d/, Varnet *et al*. 2013; /d/-/g/, Varnet *et al*. 2015a; /a/-/i/ Brimijoin *et al*. 2013), but also for different groups of listeners (Varnet *et al*., 2019, 2016, 2015b), and different maskers (white noise, Varnet *et al*. 2013; speech-shaped noise, Osses and Varnet 2021; bubble noise, Mandel *et al*. 2016).

The above set of findings based on microscopic approaches (frozen noise or ACI) confirms that EM and MM effects must be further complemented by the assessment of a token-specific effect, although the size of this effect remains an open question. For example, Régnier and Allen (2008) measured the variance caused by the masker in an auditory representation of the phoneme /t/ embedded in white and speech-shaped noises, concluding that the SNR range in which noise and speech information interacted was in fact very limited and that different noise tokens should not significantly impact phonetic judgments. Nevertheless, the T-F representations or “AI grams” adopted by Régnier and Allen are based on the concept of EM, and therefore they actually did not consider the potential effect of random envelope fluctuations into their analyses. More generally, token-specific effects are usually considered negligible by researchers in the case of stationary maskers, probably because two tokens of a steady noise generated by the same statistical process are often considered as perceptually indistinguishable. However, there is evidence that this is not generally true, as particular tokens of white noise can be discriminated (Goossens *et al*., 2009) and memorized *(Agus et al*., 2009; Pfafflin, 1968).

Below, we discuss two research areas where, from a different perspective, token-specific effects on speech perception have already been considered. The first example is related to simulations of listening performance using human-inspired auditory models. These models are composed of a front-end that accounts for the sound processing between the outer ear and the early stages of brain processing (Osses *et al*., 2022) followed by a decision back-end module that selects a response among a set of predefined alternatives (e.g., Jürgens and Brand, 2009; Zaar and Dau, 2017). This decision back end often compares processed internal representations with prototypical representations (templates) of the possible answers executed on a trial-by-trial basis. The resulting simulation data are often analyzed macroscopically, e.g., to predict overall SRT in different listening situations (Holube and Kollmeier, 1996; Steinmetzger *et al*., 2019), they may also be analyzed microscopically to examine, e.g., patterns of phoneme confusions (Jürgens and Brand, 2009). These models are particularly sensitive to the variability in the stimuli (e.g., the noise maskers), causing variability in the model internal representations. Thus, auditory models of this type are sensitive to token-specific information.

The second example where token-specific effects have received considerable attention is related to dip listening (Cooke, 2006). Dip listening refers to a listeners ability to catch brief glimpses of a speech target when the level of the fluctuating masker momentarily drops. The dip listening benefit, sometimes referred to as masking release, depends critically on the exact T-F location of the glimpses and this varies from trial to trial (Mandel *et al*., 2016). However, stationary or quasistationary noise maskers, such as the ones investigated in the present study, produce a very limited number of effective glimpses, thus allowing only a marginal dip-listening effect (Cooke, 2006).

### C. Objectives and preregistered hypotheses

The present study investigated the token-specific effect of random envelope fluctuations introduced by noise^1^ on the perception of speech acoustic cues in a consonantin-noise discrimination task using nonsense words of the structure vowel-consonant-vowel (VCV). More specifically, the task is tested using the utterances /aba/ and /ada/ presented in three types of background noises. All noises have the same flat long-term averaged spectrum but differ in the amount of random envelope fluctuations: A Gaussian white noise, a noise with a band-limited modulation spectrum, or a noise with randomly-imposed bursts of energy. The overall effect of noise on the participants’ performance is quantified from the SNR required to successfully discriminate the target sounds with a 70.7% correct response rate. Additionally, the token-specific effect of noise on performance is quantified from an analysis of the exact trial-by-trial random envelope fluctuations that is inherent to the reverse-correlation method used to derive ACIs.

In this context, our working hypotheses (H1–H4) are:

***H1***: Using the derived ACIs and the specific set of noises used during the experiments, we can predict the response (“aba” or “ada”) of each participant with an accuracy that is significantly above chance. The prediction performance metrics will provide us with a measure of the strength of the token-specific effect in this speech-in-noise task. Our expectation is that the effect should be small (Zaar and Dau, 2015) but large enough to yield predictions significantly above chance.
***H2***: Noise conditions differing only with respect to their modulation content will induce a different non-EM effect. More specifically, for a given overall performance level, noises with a larger amount of random envelope fluctuations will yield a higher ACI prediction performance. We will investigate this hypothesis by comparing the ACIs derived from three noise conditions at their corresponding (overall) SNR thresholds.
***H3***: The ACIs will be globally similar for all individuals and conditions. Not only the token-specific effect should be measurable in each listener, but it should impact the same cues for every participant, as the individual listening strategies should be globally similar. This will be confirmed by measuring the correlation between individual ACIs and the cross predictions in within-noise and between-noise conditions. If indeed the ACIs between participants are sufficiently similar, we will be able to use the ACI from one participant/condition to predict the responses of another participant/condition with significantly better-than-chance accuracy.
***H4***: Noises with a larger amount of random envelope fluctuations should yield better predictions when ACIs are derived using simulated responses from an artificial listener. This hypothesis is used to check the underlying assumptions of using ACI prediction performance as a proxy for the magnitude of the token-specific effect. The artificial listener is a perceptually-based model of the human auditory system, that adopts a fixed strategy to perform the /aba/-/ada/ discrimination. We chose a model framework that is based on the work by Dau *et al*. (1997a) and has been shown successful in simulating the overall effect of noise for a number of classical psychoacoustic tasks (e.g., Dau *et al*., 1997b; Jepsen *et al*., 2008; Osses, 2018; Osses and Kohlrausch, 2021). Given that the artificial listener strongly relies on signal-driven (bottom-up) cues using a fixed decision strategy, we will show that the specific set of noises used by the different participants has no effect on the shape of the ACI.

All hypotheses (H1–H4) were preregistered before data collection (osf.io/4ju3f/).

## II. MATERIALS AND METHODS

All stimuli and procedures described in this section were preregistered (osf.io/4ju3f/) and can be reproduced with the fastACI toolbox (Osses and Varnet, 2022b), which in the following we refer to as “the toolbox.”

### A. Apparatus and procedure

The experiments were conducted in one of the two doubled-walled soundproof booths at ENS. The experiment utilized a within-subject design. In each trial, the words /aba/ or /ada/ were presented diotically via Sennheiser HD 650 circumaural headphones (Sennheiser, Wedemark, Germany) in one of three background noises. The task of the participant was to indicate whether they heard the word “aba” or “ada” by pressing “1” or “2” on the computer keyboard, respectively.

#### 1. Experimental protocol

The participants completed 4000 trials using each noise type (total of 12000 trials), organized in thirty test blocks of 400 trials. The evaluation of a block took between 12 and 15 minutes. The order of the test blocks was pseudo-randomized with the only constraint that the three noise conditions were presented in permuted order in the first three blocks. Subsequently, all blocks were randomly assigned. Each participant required five or six two-hours sessions to complete the experiment.

For each test block only one type of noise was evaluated and one independent adaptive track was measured: After a correct or incorrect response, the level of the target word in the subsequent trial was decreased or increased, respectively, following a one-up one-down weighted adaptation rule (Kaernbach, 1991). We used up- and down-steps in a ratio of 2.41 to 1 that lead to a target a score 70.7% according to Kaernbach (1991, his Eq. 1). Participants received feedback on the correctness of the trial. Furthermore, they were explicitly instructed to minimize their response bias as much as possible with a warning message displayed on screen when the response ratio was higher than 60% or lower than 40%.

For each trial, we stored the participant’s response, the corresponding SNR, the target actually presented, and the exact waveform of the noise. After completing each block, participants were encouraged to take a short break.

#### 2. Training session

Before the first test block, the participants completed a short training to make sure that they correctly understood the task. This training session was similar to the main experiment except that participants were able to repeat the noisy speech stimuli or to listen to /aba/ or /ada/ samples in silence. The training ended when the participant was ready to start the main experiment. The training results were excluded from any further analysis.

### B. Participants

Twelve participants (S01–S12) aged between 22 and 43 years old (4 females, 8 males) took part in our study, with eight of them being native French speakers. This information is presented in Appendix A1. The participants provided their written informed consent prior to the data collection and were paid for their contribution.

The participants’ hearing status was measured using pure-tone audiometry at six frequencies (250, 500, 1000, 2000, 4000, and 8000 Hz) and had average thresholds of 20 dB HL or better in their best ear. The obtained hearing thresholds are shown in Fig. A.1.

The total number of *N* = 12 participants exceeds the preregistered number of participants (*N* = 10). By the time we completed the data collection for *N* = 10, twelve participants had been recruited, and we therefore decided not to interrupt the data collection for the last two subjects (S06 and S12). The results reported in this study are based on all twelve participants. The same analysis using the preregistered sample size can be found in Appendix A3 b and yielded very similar results.

### C. Simulated participant: The artificial listener

In addition to the experimental data collection, we used an auditory model to simulate the performance of an average normal-hearing listener who uses a fixed decision criterion to compare sounds. This “artificial listener” assesses the internal representations of each sound using signal-driven (bottom-up) information based on a modulation-filter-bank approach (Dau et al., 1997a). The internal representations were subsequently compared using a (top-down) decision back-end based on template matching, with two stored templates, one for each target sound (Osses and Kohlrausch, 2021). The artificial listener was treated as an additional participant, meaning that its results were subjected to the same data analysis as applied to the experimental data.

The auditory model is composed of a front-end and a back-end processing using default parameters in our toolbox. In short, the front-end processing is very similar to the model described by Osses and Kohlrausch (2021), except that the middle-ear module is implemented as a linear phase filter and that the modulation filter bank uses a Q factor of 1. The two templates were derived at a supra-threshold SNR of −6 dB where each target was embedded and subsequently averaged across 100 newly-generated white noise realizations. This fixed white-noise template was used for the simulations in all three noise conditions. The trial-by-trial decision was based on a template-matching where a decision bias was introduced to allow the model to balance the number of “aba” and “ada” choices (Osses and Varnet, 2021). All details about the model configuration and the decision scheme can be found in Appendix A4.

### D. Stimuli

#### 1. Target sounds

We used two male speech utterances from speaker S43M taken from the OLLO speech corpus (Meyer *et al*., 2010) (words /aba/: S43M_L007_V6_M1_N2.wav; /ada/: S43M_L001_V6_M1_N1.wav). We preprocessed these speech samples to align the time position of the vowel-consonant transitions, to equalize their energy per syllable, and to have the same waveform duration. The stored sounds have a duration of 0.86 s, a sampling frequency *f_s_* of 16 kHz, and an overall level of 65 dB sound pressure level (SPL). The time-frequency (T-F) representation of the stored sounds is presented in Fig. 1, together with their fundamental-frequency *(f_0_)* and formant (*F*_1_–*F*_4_) trajectories.

Prior to the stimulus presentation and the addition of noise, the sounds were adjusted in level depending on the corresponding SNR.

#### 2. Background noises

Three types of background noise conditions were tested. These conditions were chosen to include a stationary noise condition using white noises and two additional noises with stronger envelope fluctuations below 35 Hz, that correspond to non-stationary noise conditions. To increase the low-frequency fluctuations in the envelope domain, we designed an algorithm to generate a white noise with superimposed Gaussian bumps and an algorithm to generate noises with limited modulation power spectrum. We abbreviate these two types of noises as bump and MPS noises, respectively. The noises were generated at an *f_s_* of 16 kHz, were set to have an overall level of 65 dB SPL and were subsequently gated on and off with 75-ms raised-cosine ramps before being stored on disk.

The acoustic characteristics of the noises are shown in Fig. 2 and the algorithm details are given in the next paragraphs. For each noise we show the T-F representation of one arbitrarily-chosen noise realization (Fig. 2**A**), followed by an acoustic analysis of the noises derived from 1000 noise realizations (Fig. 2**B**–**C**) using (1) critical-band levels within 1 ERB for bands centered between 87 Hz (or 3 ERB_N_) and 7819 Hz (or 33 ERB_N_) in Fig. 2**B**; and (3) assessing the broadband envelope spectrum obtained from the absolute value of the Hilbert envelope, downsampled to *f_s,env_* = 1000 Hz. The envelope spectrum after DC removal is shown in Fig. 2**C**^2^.

**FIG. 2.**
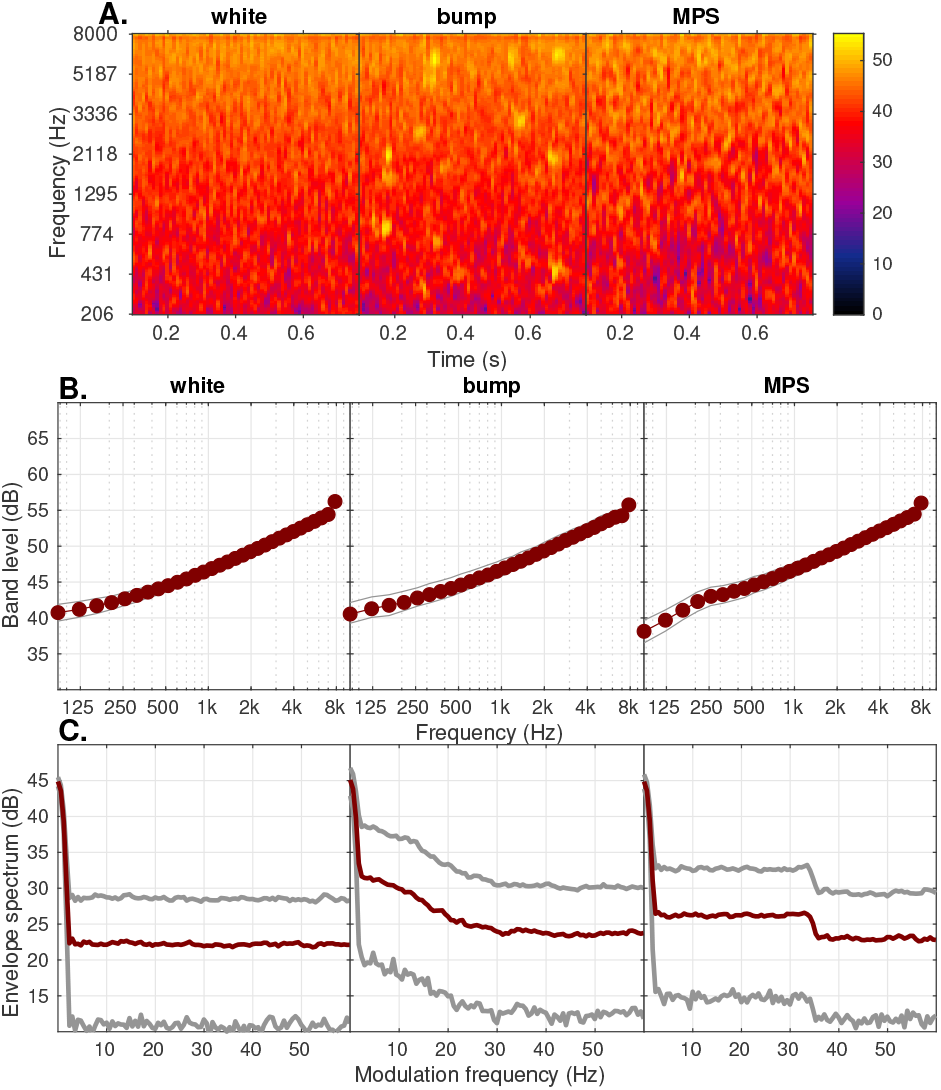
Summary of the acoustic characteristics of white (left), bump (middle), and MPS noises (right). **A**. T-F representation of one arbitrary-chosen noise representation. **B**. Critical-band levels within 1-ERB wide filters. **C**. Envelope (modulation) spectrum. More details are given in the text. In Panels **B**–**C**, the gray curves indicate the percentiles 5 and 95 of the corresponding estimate.

The generated white noises had a spectrum level of 26 dB/Hz with an effective bandwidth between 0 and *f_s_*/2, resulting in critical-band levels between 40.7 dB and 56.2 dB (Fig. 2**B**). The envelope spectrum (Fig. 2**C**) was approximately constant with a median amplitude of 22.2 dB, although a theoretical monotonic decrease up to *f_s_/2* is expected (Dau *et al*., 1999). This decrease is not visible due to the 0–60 Hz limit of the abscissa.

The bump noises were generated using an algorithm similar to that described by Varnet *et al*. (2019). The bumps are regions of excitation that have a Gaussian shape defined by a temporal width of *σ_t_* = 0.02 sec and a spectral width of *σ_f_* = 0.5 ERB, emphasized up to 10 dB. The time and frequency locations of the bumps were randomly spread across the entire duration of the waveform and through the whole T-F space, i.e., between 80 Hz and 7158 Hz (i.e., 1 ERB_N_ below 8000 Hz). Each waveform contained 30 newly drawn Gaussian bumps. The generated bump noises had critical-band levels between 40.5 and 55.8 dB (Fig. 2**B**). The envelope spectrum (Fig. 2**C**) had a triangular shape going from an amplitude of 31.4 dB at *f*_mod_ = 3 Hz down to 23.5 dB at *f*_mod_ = 31.1 Hz with an approximately constant spectrum thereafter (median amplitude of 23.7 Hz).

Finally, the MPS noises were generated by limiting their spectrum in the modulation frequency domain using a set of temporal and spectral rate cut-off frequencies. We chose a temporal cut-off of 35 Hz and a spectral cut-off of 10 cycles/Hz, based on the study by Elliott and Theunissen (2009) and some pilot tests. The MPS bandwidth was limited using the phase reconstruction approach from the PhaseRet toolbox (Průŝa, 2017). Inspired by Venezia *et al*. (2016), we first generated a white noise that is multiplied in the MPS domain by a low-pass envelope with the desired characteristics defined by the temporal and spectral cut-off frequencies. The MPS-limited representation is then converted back to a time-domain waveform and stored on disk. The generated MPS noises had critical-band levels between 38.1 and 56.0 dB (Fig. 2**B**). The envelope spectrum (Fig. 2**C**) had a constant value of 26.3 dB starting after the DC and up to about 35 Hz, an amplitude that decreases to a constant value of 23.0 Hz thereafter.

In summary, all noises originate from white noises with or without emphasized envelope fluctuations, sharing nearly the same long-term spectral content (Fig. 2**B**). However, the noises have a different amount and distribution of random envelope fluctuations in the modulation-frequency domain (Fig. 2**C**). White noises have low envelope fluctuations, MPS noises have a rectangular-shaped low-pass envelope fluctuations (cut-off *f* _mod_ =35 Hz), and bump noises have a triangular-shaped low-pass envelope fluctuations (for *f*_mod_<31.1 Hz).

For each participant a new set of 4000 noises with a level of 65 dB SPL was generated, resulting in 36 sets of noises (12 participants ×3 noise conditions). The simulated participant, the artificial listener (Sec. IIC), was tested on the same 36 sets of experimental noises. The waveforms in each noise were stored and can either be retrieved from Zenodo (Osses and Varnet, 2022c) or be reconstructed using the toolbox (see Appendix A2).

#### 3. Noisy trials

In each trial, the level-adjusted target sounds were arithmetically added to the corresponding noise according to the adopted staircase rule. Before the trial was administered to the listeners, an additional but small variation in the total presentation level (level roving) between —2.5 and +2.5 dB (continuous range, uniform distribution) was applied to partly discourage the use of loudness cues during the experiment. For the simulations using the artificial listener that were run with all 36 noise data sets, the same order of trials and level roving were applied as used for each participant.

### E. Data analysis

The experimental data collection resulted in 36 staircases for twelve participants in the three noise conditions and 36 staircases for the simulations with the artificial listener. In this section we describe the analyses that were applied to the experimental trials to obtain the direct behavioral results and to derive the individual ACIs.

#### 1. Preregistered data exclusion criteria

For a more efficient data processing, the ACI method requires a minimization of response biases. Next to the explicit instruction to balance “aba” and “ada” choices (Sec. IIA 1), we preregistered two criteria for trial exclusion.

The first criterion is related to the exclusion of all starting trials of each test block before reaching the fourth turning point or reversal. Those trials correspond to the so-called approaching phase of the staircase procedure, where the adjustable parameter, the SNR, is considered to be at a supra-threshold level with a percentage correct that is well above the target 70.7%. The fourth reversal was considered to be the staring point of the measuring phase of the staircase.

The second criterion is an explicit control of the balance between “aba” and “ada” responses in our dataset. During the data processing, the responses of the target sound that obtained more preferences were sorted in increasing SNR. Subsequently the trials with most extreme values (minima or maxima) were discarded until the same number of “aba”-”ada” preferences was achieved. In other words, if a participant indicated “aba” 53% of the times and “ada” 47% of the times, the trial exclusion was applied to the “aba” trials.

#### 2. Measures of behavioral performance

The listeners’ performance in the different noise conditions was assessed using a number of measures derived from the trial SNRs. The percentage of correct responses and SNR thresholds were obtained for each block of 400 trials, after data exclusion (Sec. IIE 1). Then, the rate of correct responses in /aba/-trials and in /ada/-trials were expressed in histograms using SNR bins of 1 dB. Note that these values correspond to the rate of hit and correct rejection if we arbitrarily identify /aba/- and /ada/-trials as target-present and target-absent trials, respectively. Finally, using the same 1-dB wide bins, the classical discriminability index (*d*′) and criterion (c) metrics were obtained from the hit, false alarm, correct rejection, and miss rates (Harvey, 2004) as a function of SNR.

The behavioral measures *d′*, c, and the block-by-block SNR threshold were tested for a group-level effect using a mixed analysis of variance (ANOVA) with two fixed factors, block number and noise condition. Participants were treated as a random effect, meaning that differences in baseline performance for individual listeners were taken into account. A second mixed ANOVA with two fixed factors, SNR and condition, was run to confirm the effect of SNR on *d*′. Similarly, a mixed ANOVA was also run on the criterion *c*.

#### 3. Auditory classification images (ACIs)

##### a. Time-frequency (T-F) representations

Following the same rationale as in previous studies, the ACIs were derived and interpreted in a T-F space (Osses and Varnet, 2021; Varnet *et al*., 2013, 2015a). Here, we chose to use a Gammatone-based representation rather than a spectrogram. The 0.86-s long monaural noises were decomposed into 64 bands equally spaced in the ERB-number (ERB_N_) scale (Glasberg and Moore, 1990) between 45.8 Hz (1.69 ERB_N_) and 8000 Hz (33.19 ERB_N_), spaced at 0.5 ERB. The filters had a width of 1 ERB, resulting in a 50% overlap. The 64 band-passed signals were then low-pass filtered using a Butterworth filter (f_cut-off_=770 Hz, fifth order), which roughly simulates the inner-hair-cell envelope extraction processing (see, e.g., Osses *et al*., 2022, their Sec. 2.4). Finally, one estimate every 0.01 s (amplitude mean) was obtained for each of the frequency bands along the time dimension resulting in a final T-F noise representation stored in a 86-by-64 matrix. We denote the T-F representation of the noise presented to participant *k* in trial *i* as 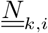, while *<u>N</u>_k,i_* refers to the vectorization of this matrix. In the following sections, we use the same formalism to refer to the ACI in its matrix form (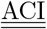, 86-by-64) or vector form (<u>ACI</u>, 5504-by-1).

##### b. Generalized linear model

The core principle of the ACI approach is to assess how the random envelope fluctuations in the stimulus 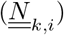 affect the behavioral response of the participant (denoted *r_k,i_*) on a trial-by-trial basis. For this purpose, we relied on a stimulus-response transformation based on a generalized linear model (GLM) to produce a T-F matrix of decision weights (Varnet *et al*., 2013, 2015a). As the objective of our study was to isolate the effect of random envelope fluctuations on phoneme perception, the GLM did not include any complementary predictor like the target actually presented or the SNR. We define the vectorized ACI for participant *k*, <u>ACI</u>_*k*_, such that:

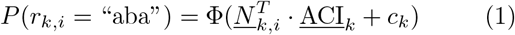

where *P*(*r_k,i_* = “aba”) = 1 – *P*(*r_k,i_* = “ada”) is the predicted probability of choosing “aba” and Φ(*x*) stands for the sigmoid function Φ(*x*) = 1/ (1 + *e*^-*x*^). Equation1relates the specific content of the noise in trial *i* to the response given by the participant, with <u>ACI</u>_k_ and *c_k_* being the GLM parameters that need to be fitted to each participant’s data. The <u>ACI</u>_*k*_ in Eq. 1 is expressed as a vector of perceptual weights, with each element corresponding to one T-F point of the noise representation *<u>N</u>_k,i_*. There-fore, they are more easily interpreted as a matrix 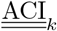 that has the same size as the 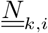 matrix. The parameter *c_k_* corresponds to the fitted intercept value that indicates the overall bias of the participant towards one response or the other.

##### c. Sparseness prior in a Gaussian pyramid basis

An ACI is typically composed of many non-zero weights. However, these weights are often grouped into positive or negative clusters matching the location of acoustic cues in the targets, while the rest of the T-F space is close to zero. Therefore, a more compact way of describing an ACI would be as a linear combination of Gaussian-shaped elements centered at different T-F locations, such as the ones shown in Fig. 3. Here, formulating the problem in a space where ACIs can be expressed with a limited number of coefficients allows us to enforce “sparse” solutions, that is, ACIs that are non-zero only in a few localized T-F regions.

**FIG. 3.**
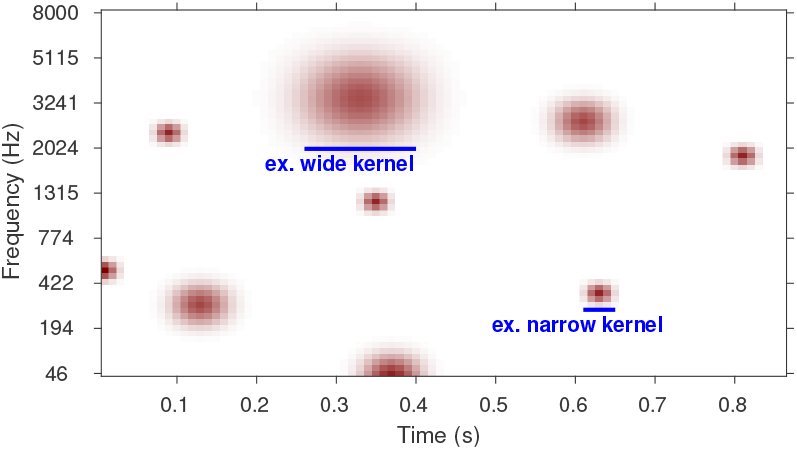
Illustrative example of a nine Gaussian basis element used in the Pyramid decomposition represented in a T-F space, as used for the ACIs. Thanks to the multi-resolution matrices of the decomposition method, narrower and wider cues can be extracted from the input noise matrix *S_k,i_*. Two stereotypical basis elements are indicated by the blue lines.

The Gaussian pyramid consists of four successive levels (1 to 4) corresponding to decreasing T-F resolutions. Each level is composed of Gaussian elements of the same width (standard deviation = 1, 2, 3, or 4 bins, respectively) and spaced every 1, 2, 3, or 4 bins. This means that the first level is not subsampled in contrast to the gradually more subsampled levels 2 to 4.^3^ The coefficients of the Gaussian elements from all four levels are normalized to have a norm equal to 1, vectorized, and stored into a single matrix 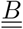. An ACI is then expressed as a linear combination of Gaussian elements:

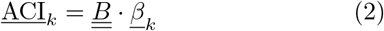

which represents a change of basis that relates the coordinates of the ACI in the new multi-resolution Gaussian-pyramid space *(<u>β</u>_k_*), to its coordinates in the T-F space. By replacing <u>ACI</u>_*k*_ in Eq.1 we obtain:

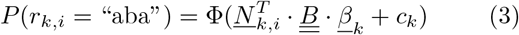

Instead of estimating the ACI directly in the T-F space, using Eq. 1, we actually estimated the <u>*β*</u>_*k*_ coefficients of Eq. 3. Although Eqs. 1 and 3 are mathematically equivalent, the latter allows the prior assumption about the simplicity of the ACI to be expressed by looking for as few non-zero *β_k_* coefficients as possible (Mineault *et al*., 2009). In statistical terms, this is achieved by penalizing the classic maximum-likelihood estimator with a L1-regularized (lasso) regression approach, which enforces sparse solutions. The weight and bias for each participant (<u>*β*</u>_*k*_ and *c_k_*) were fitted individually with a lasso regression, then transformed back into the T-F space using Eq. 2 to obtain the final 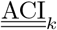. Following the standard lasso procedure, the hyperparameter λ that controls the strength of the regularization was selected to minimize the deviance through a 10-fold crossvalidation approach. We tested twenty plausible λ values logarithmically spaced between 1.1 ·10^-3^ and 0.1, with larger values enforcing more sparse candidates, as shown in Fig. A.7 (see also Varnet et al., 2015a, their Fig. 2). The choice of this range of values ensured that the lowest amount of regularization was low enough to produce a very noisy ACI, while the highest amount of regularization was high enough to produce a flat ACI, as shown in the left-most and right-most panels of Fig. A.7, respectively. A flat or “null” ACI, only contains weights that are equal to zero, meaning that an ACI prediction is only defined by the bias *c_k_*. This statistical-fitting procedure is the same as we have used in our last studies (Osses and Varnet, 2021, 2022a).

#### 4. Out-of-sample prediction

##### a. Performance metrics

Following the standard hyperperameter selection procedure for GLMs (e.g., Mineault et al., 2009; Varnet et al., 2015a;Wood, 2017), the out-of-sample predictive performance of the fitted ACIs was assessed during the 10-fold cross-validation using the cross-validated deviance. To allow a direct comparison between different ACIs, that differ in the exact number of test trials due to the criteria for trial exclusion (see Sec. II E 1), we report the cross-validated deviance per trial (CVD_t_).

We adopted a second complementary metric that we defined as prediction accuracy (PA). PA is a “noisier” but more intuitive measure of prediction performance that is assessed as the coincidence between predicted and actual responses. PA relates the predicted and actual responses, expressing “aba” (or “ada”) predictions when /aba/ (or /ada/) was actually chosen by the participant. Assuming that a probability *P* equal to or above 0.5 in Eq. 1 (or Eq. 3) would be related to a predicted choice of “aba”, the PA metric can be formalized as:

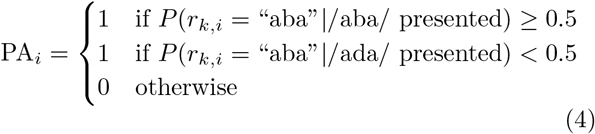

This metric was summed across trials and expressed as a percentage and can adopt values between chance (~50%) and 100%.

To facilitate the interpretation of the previous performance metrics, we define the deviance-per-trial benefit ΔCVD_t_ and the percent accuracy benefit ΔPA as the difference between prediction performance using the optimal ACI (Eq. 3) and that of the corresponding null ACI. For ΔPA we further scaled the metric by 1/(1-PA_null_). This way, PA values that can range between ~50 and 100% are mapped to ΔPA values between 0 and 100%.

Given that lower ΔCVD_t_ values indicate better predictions, for individual evaluations we provide one-sided 95% confidence intervals, obtained as 1.64 times the standard error of the mean (SEM), reporting a significant benefit if the confidence interval is below zero. For the group evaluation, mean ΔCVD_t_ values across folds were obtained for each participant and the significance was assessed at the group level following a similar criterion.

With the ΔPA metric, the benefit of using the optimal ACI with respect to the null ACI is expected to increase. As a reference, we show the boundaries at 2.6% or 4.78% above chance for evaluations using all experimental trials or incorrect trials only, respectively.^4^

For the purposes of this study, the metrics of prediction accuracy are not only a way to validate ACIs, but they also provide a proxy for the size of the token-specific effect on phoneme perception with better values when more of the participant confusions are due to random envelope fluctuations. Strictly speaking, the predictability gives us a lower boundary of the token-specific effect, as responses that are correctly predicted using a model that by design is only based on random envelope fluctuations—as it is the case here for the T-F representations transformed using the fitted GLMs—must be caused by those random envelope fluctuations. On the contrary, incorrect predictions could either be due to token-specific effects that are not accounted for in our GLM approach, such as interactions between separate T-F regions or effects where non-linear processes are involved, or to other causes. For these reasons, we also report the performance metrics only using data from incorrect trials. In this case, the metrics are labeled as ΔCVD_t,inc_ and ΔPA_inc_, respectively.

##### b. Cross predictions

The selected performance metrics measure the ability of the ACI, fitted on a subset of the participant’s data, to predict unseen data from the same participant in the same condition. Complementary to these “auto predictions,” we also derived “cross predictions” where the fitted ACI was evaluated on a test set extracted from a different participant or a different condition. We assessed two types of cross predictions: (1) within participant but between conditions, and (2) between participants but within conditions. While the auto predictions were used to assess the goodness-of-fit of the obtained ACIs, the cross predictions were used to evaluate the similarity between listening strategies across participants or across maskers. To test the significance of the cross predictions, two ACIs were considered as “similar” if we could exchange them and reach a significantly better-than-chance prediction accuracy, i.e., with newly obtained ΔCVD_t_ being significant according to the criteria defined above. We also report the correlation across ACIs, but this complementary analysis is only presented in Appendix A 3.

## III. RESULTS

For each participant, the experimental data collection was completed across different days, mostly requiring five two-hour sessions. All the recruited participants (*N* = 12) were able to complete the task, although participant S10 showed highly variable results with scores that were clearly below the group average.

### A. Measures of behavioral performance

In the course of the experiment, the level of the speech target (/aba/ or /ada/) was adapted using a one-up one-down weighted adaptation rule that targeted a 70.7% of correct responses (see Sec. IIA 1). In practice, after excluding the approaching phase of the staircases (see Sec. IIE 1), the exact percentage of correct responses averaged across noise conditions and test blocks ranged between 71.0% (S10) and 71.6% (S03), with session by session scores between 69.5% and 74.2%.

To provide an overview of the participants’ performance, the obtained SNR thresholds as a function of test block are shown in Fig. 4. In this figure, we show the overall performance (averaged across participants) for white-(blue), bump-(red), and MPS-noise conditions (green). Additionally, the SNRs for each participant were averaged across noise conditions obtaining twelve gray traces, also shown in Fig. 4. The thresholds of participants S04 and S10 are shown in solid lines and correspond to the (overall) best and worst performing participants in the task, respectively. The SNR evolution given by the gray traces suggests that there was a small learning effect during the course of the experiment, with slightly better (lower) thresholds in the last test blocks. This small learning effect was confirmed by a two-way mixed ANOVA with the factors masker and test block. Both factors were found to have a significant effect on the obtained SNR thresholds with *F*(2,345) = 15.87, *p* < 0.001 and F(1,345) = 36.63, *p* < 0.001, respectively. A post-hoc analysis revealed that the effect of masker type was in fact due to a difference in the bump-noise condition compared to the other two, while SNR thresholds for white noise and MPS noise were not significantly different.

In order to measure the effect of speech level on performance, trial-by-trial responses were converted to mean scores, *d*′, and criterion values (c) as a function of SNR. These metrics are shown in Fig. 5. We then ran three two-way mixed ANOVAs, independently applied to the factors masker, SNR, and criterion. For these tests we only used the data for the SNR bins centered between −16 and −12 dB, where data for all participants in all conditions had been obtained. The ANOVAs supported a significant effect of the factors masker and SNR on *d*′ (masker: *F*(2,165) = 10.39, *p* < 0.001; SNR: *F*(1,165) = 1017.65, *p* < 0.001), while only the factor SNR had a significant effect on criterion (masker: *F*(2,165) = 1.75, *p* = 0.178; SNR: *F*(1,165) = 9.77, *p* = 0.002). According to a post-hoc test, the effect of masker type on *d*′ was due to a difference in the white-noise condition compared to the other two.

**FIG. 4.**
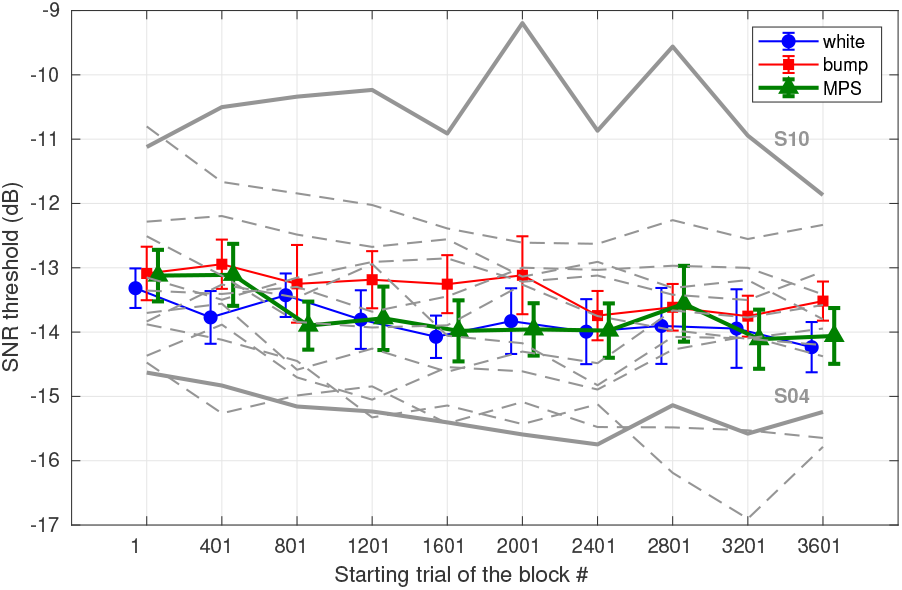
(Color online) Mean SNR thresholds for the group in blocks of 400 trials for each of the three masker conditions. We show the individual thresholds for the twelve participants averaged across conditions (gray dashed lines), emphasizing the overall thresholds of the participants with lowest and highest values (thick gray continuous lines). The error bars indicate one standard error of the mean (SEM).

**FIG. 5.**
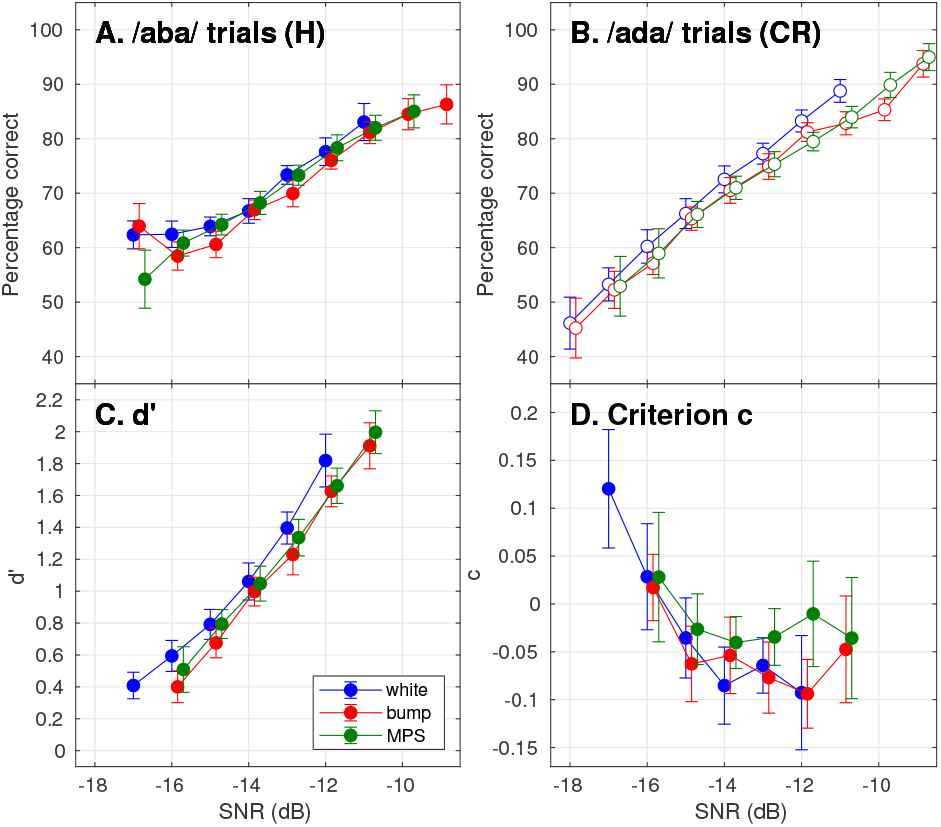
(Color online) Percentage of correct responses for (**A**) /aba/ and (**B**) /ada/ trials, (**C**) discriminability index *d′*, and (**D**) criterion, as a function of 1-dB wide SNR bins. These metrics were obtained using the group data. Error bars indicate 1 SEM.

### B. ACIs

The ACIs that were derived from the collected data are shown in Fig. 6 using white, bump, and MPS noise maskers. Panels **A–F** contain the individual ACIs, while a group average is shown in the bottom-most panels (Fig. 6**G–I**). Overall, the individual ACIs bear some similarities, although for two participants (S09 and S10) the hyperparameter selection in white noise did not yield a minimum, resulting in a null ACI (dashed pink boxes in Fig. 6). Large weights were found at *t* ≈0.3 s, the time of the onset of the second syllable, which are more clearly visible in the group ACIs. More specifically, we found a clear pattern of positive (red) and negative (blue) weights matching the location of the *F*_1_ and *F*_2_ onsets of the /aba/ and /ada/ sounds. Additionally, in a subset of ACIs, weak but consistent perceptual weights were also found around the time of the first-syllable *F*_2_ offset (e.g., in the ACIs of S03, all conditions), or near the release burst of the plosive consonant at around 8 kHz (e.g., ACI of S07, white-noise condition).

**FIG. 6.**
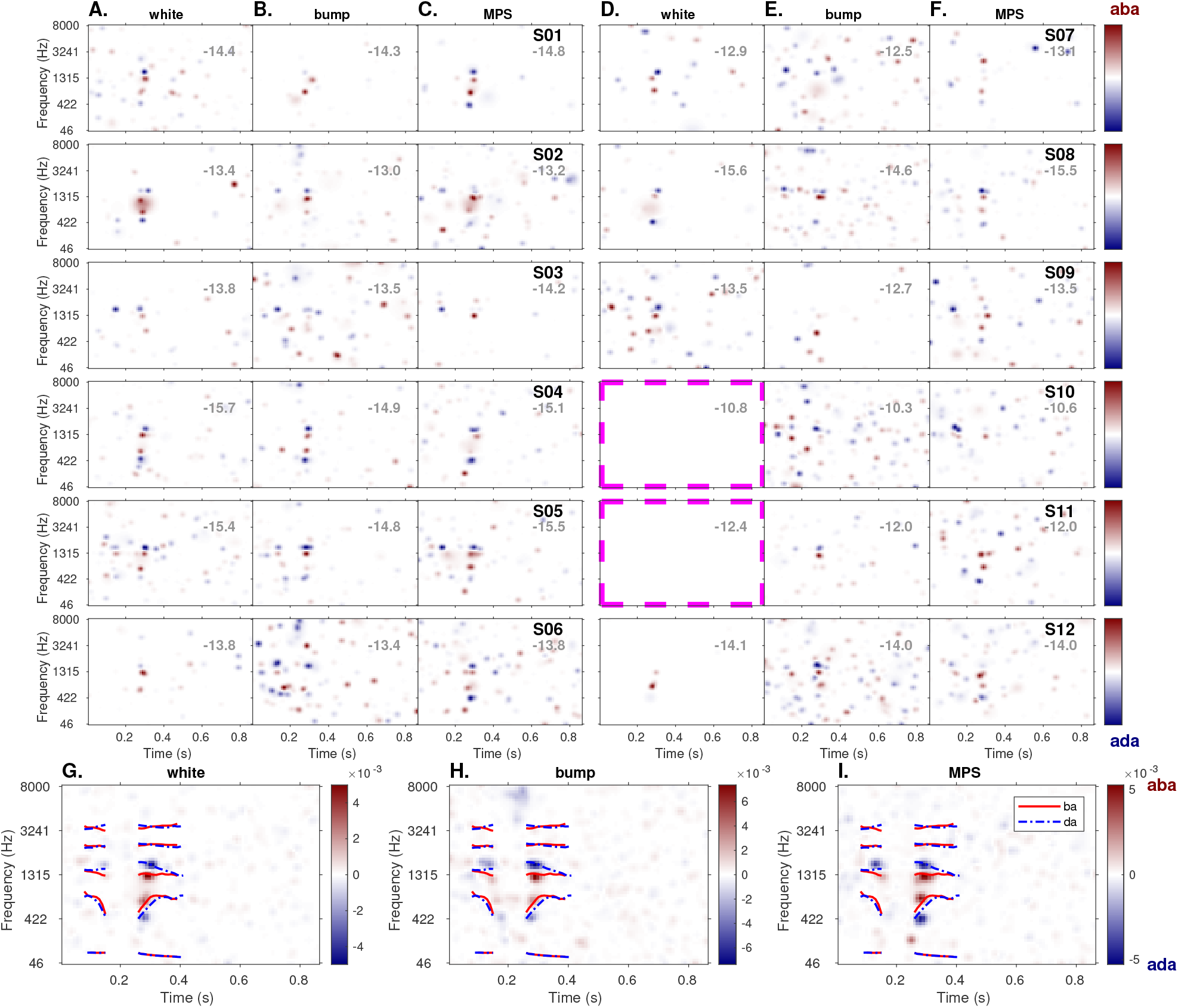
(Color online) Top panels: ACIs for the 12 participants using (**A,D**) white, (**B,E**) bump, and (**C,F**) MPS noises. For comparison purposes, the weights in each ACI are normalized to their maximum absolute value. The values in gray in the top right corner of each ACI indicate the corresponding mean SNR threshold expressed in dB. The dashed pink boxes indicate the two ACIs that only contain zero weights. Bottom-most panels (**G**–**I**): mean ACI across all participants, in each condition. The formant trajectories for /aba/ (red solid lines) and /ada/ (blue dotted lines) are superimposed.

The group ACIs were obtained as the arithmetic average of all non-normalized individual ACIs for white (Fig. 6**G**), bump (Fig. 6**H**), and MPS noises (Fig. 6**I**) and are only shown for visualization purposes. In these panels we superimposed the *f*_0_ and formant trajectories of /aba/ and /ada/. The group ACIs were not normalized to emphasize the fact that the (blue and red) weights have different limits for the different noises.

### C. Out-of-sample prediction accuracy

#### Auto predictions at the individual level

The out-of-sample metrics of prediction accuracy, ΔCVD_t_ and ΔPA, at the individual and group level are shown in Fig. 7, where the metrics for the individual ACIs are shown as open diamond markers.

**FIG. 7.**
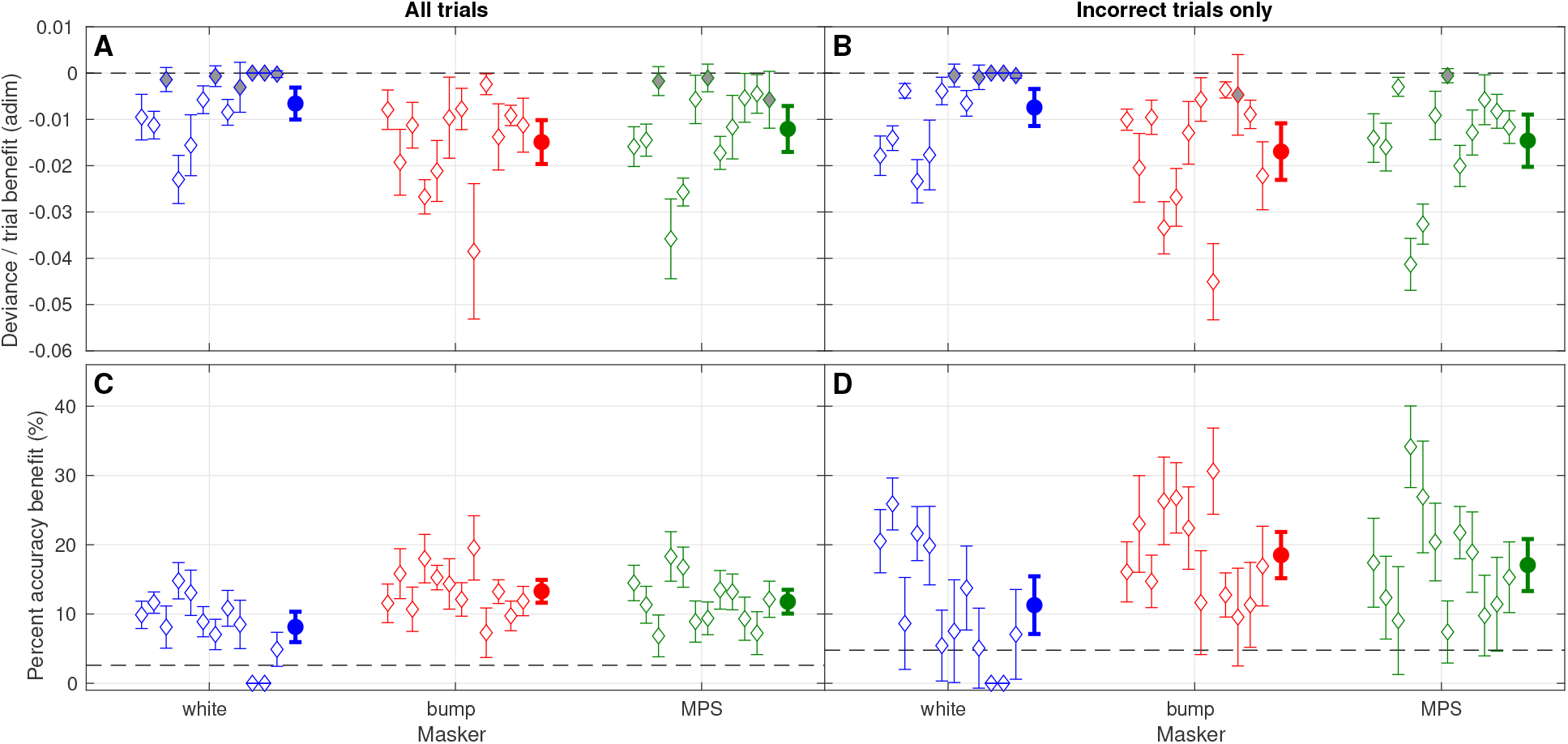
(Color online) Metrics of performance benefit, ΔCVD_t_ (top panels) and ΔPA (bottom panels), for each of the obtained ACIs, from left to right for participants S01 to S12 (open or gray diamonds). The filled circle markers indicate the group average for the corresponding condition. Left (**A, C**) and right (**B, D**) panels indicate the analysis using all trials or the incorrect trials only, respectively. In all cases, the error bars indicate ±1.64 SEM and the gray markers indicate those estimates that were found to be non significant, based on the boundaries at 0 (Panels **A** and **B**, dashed lines).

The results based on ΔCVD_t_ (Fig. 7**A**) show that 9 ACIs out of 36 did not yield predictions significantly higher than chance. Those non-significant estimates are marked in gray in Fig. 7**A**. From these 9 ACIs, two corresponded to the conditions where a null ACI was obtained (dashed pink boxes in the figure), which, by definition, are related to performance metrics equal to zero.

For the analysis of incorrect trials, the results are shown in Fig. **7B** and **7D** for ACVD_t,inc_ and APA_inc_, respectively. Although the improvement in ACVD_t,inc_ was rather small (compare Fig. **7B** with 7**A**), there was a systematic improvement of APA_inc_ values (compare Fig. **7D** with 7**A**).

#### Auto predictions at the group level

At the group level, white noise yielded a smaller prediction performance compared to the bump and MPS noise conditions with ACVD_t_ values of −0.657 · 10^-2^, −1.490 · 10^-2^, and −1.207 · 10^-2^ (filled maskers in Fig. 7**A**) and ΔPA values of 8.1, 13.3, and 11.8%, respectively (filled maskers in Fig. 7**C**). Additionally, a significant effect was found for the factor “masker” in a mixed ANOVA on ACVDt (*F*(2, 22) = 7.73, *p* = 0.003).

When restricting the test set to incorrect trials only (Fig. 7**B**, 7**D**), the prediction accuracy was found to be systematically higher: From the participants’ incorrect answers a larger proportion of these errors is explained using the ACIs in bump-(ACVD_t,inc_ = −3.390 ·10^-2^; APA_inc_ = 18.5%) and MPS-noise conditions (ACVD_t,inc_ = −2.920·10^-2^; APA_inc_ = 17.1%) compared to the whitenoise condition (ACVD_t,inc_ = −1.485·10^-2^; APA_inc_ = 11.3%). These group benefits are indicated by filled markers in Fig. **7D** and are all significantly above chance.

#### Cross predictions between participants

The procedure to obtain cross predictions between participants (see Sec. II E 4 b) resulted in three 12-by-12 matrices of cross-prediction values when the data of one participant in one noise condition was predicted using the ACI from another participant in the same condition. The obtained APA values are shown in Fig. 8**A**-**C** and ranged between −1.4% and 19.5%. The main diagonal of these matrices correspond to the same auto-prediction values that are shown as open diamond markers in Fig. 7**C**. As expected, the ΔPA values were overall lower (or overall higher using ACVDt) than the auto-prediction values, with on-diagonal averages of 8.1, 13.3, and 11.8% for white-, bump-, and MPS-noise conditions, respectively, and corresponding off-diagonal averages of 4.7, 6.0, and 5.9%. From the off-diagonal metrics and only considering the ACIs that produced significant predictions, i.e., excluding auto predictions and the cross predictions indicated by the red arrows in Fig. 8**A**-**C**, 42 (out of 66), 67 (out of 132), and 57 (out of 99) cross predictions led to a performance that was significantly above chance for white-, bump-, and MPS-noise conditions, respectively. Those cross predictions are enclosed in pink dashed boxes in the figure. With this significance analysis, we can identify the ACIs that better predict the data or the data that are better predicted by other ACIs, by looking at the vertical or horizontal direction of the corresponding matrix, respectively. For instance, the ACI from S11 in Fig. **8C** produced significant predictions using the data of two participants (S01, S05, vertical direction) and the ACI from S05 produced significant predictions using the data of all participants except one (S11). In the cross prediction of data using other ACIs, the data from S09 for white noise was significantly predicted only using the ACI from S08 (Fig. 8**A**, horizontal direction), while the data from S05 in the bump and MPS noise conditions (Fig. 8**B**, 8**C**), were significantly predicted using eight (of 11) other ACIs.

**FIG. 8.**
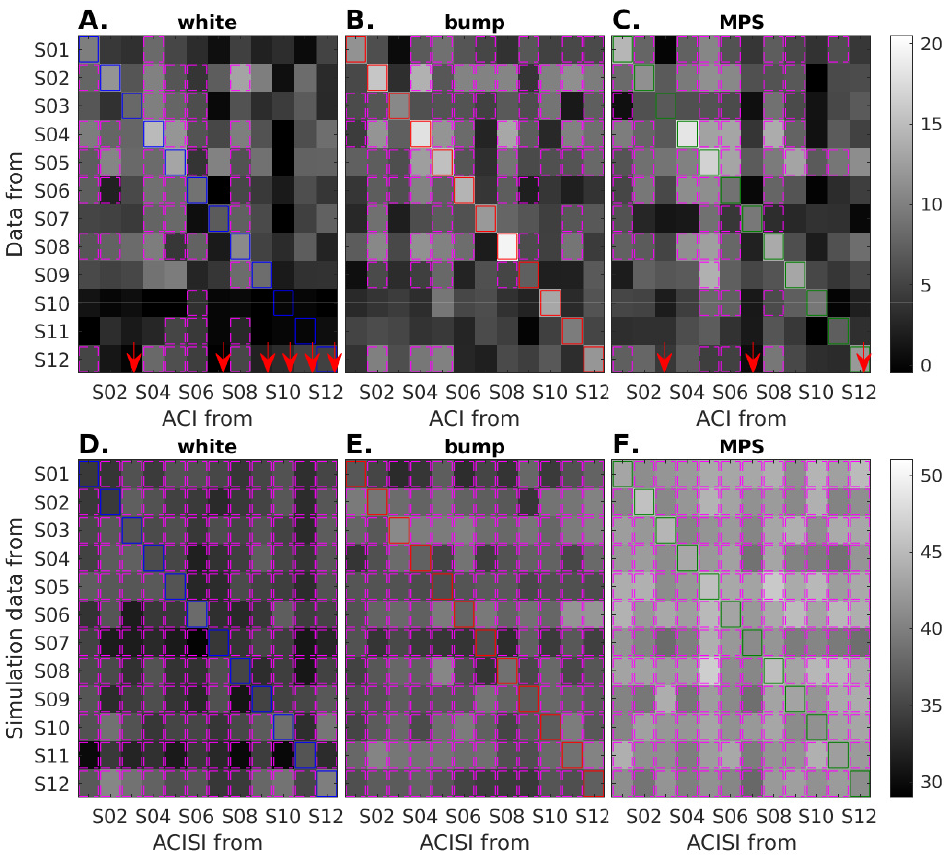
(Color online) **A** –**C**. Between-subject cross-prediction matrices for the three conditions using ΔPA, expressed as corrected-for-chance percentages. The main diagonals are enclosed in colored squares and correspond to the same auto-prediction values as in Fig. 7**C**. The pink dashed boxes indicate the ACIs from the abscissa that are able to predict significantly above chance the data of the participant indicated in the ordinate. See the text for details and also Fig. A.3. From this analysis, we excluded the ACIs that did not lead to significant auto predictions (red arrows), which were marked in gray in Fig. 7**A**. **D**–**F**. Same cross-prediction analysis but using the simulation data. In this case, all cross predictions led to significant predictions. Note the different (higher) range of ΔPA values with respect to the top panels.

#### Cross predictions between noises

The procedure to obtain cross predictions between noises but within participant (see Sec. IIE4b) resulted in twelve individual 3-by-3 matrices, that are shown in Fig. A.4**A**. We focus on the cross predictions averaged across participants and using ΔPA, which we present in Fig. 9**A**.

**FIG. 9.**
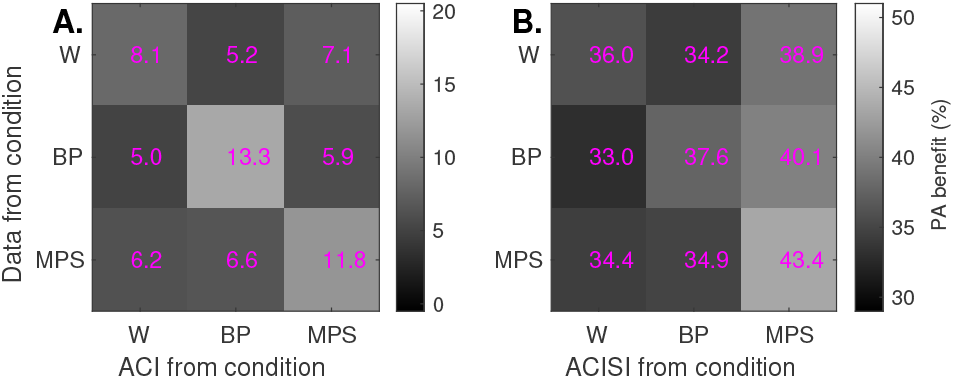
Between-noise cross predictions using ΔPA values averaged across (**A**) participants or (**B**) simulated participants. The off-diagonal values were overall lower than the within-noise predictions. See the text for further details.

The global results show that the auto (within-noise) predictions of the main diagonals gave overall ΔPA values of 8.1, 13.3, and 11.8% for the white, bump, and MPS noises (the same group values as in Fig. 7**C**) that decreased to (at most) 5.0, 5.2, and 5.9%, respectively, when using an ACI to estimate the collected data between noises (compare the elements of Fig. 9 in the vertical direction). Along the horizontal direction, i.e., when exchanged ACIs are used to predict the data within noise condition, the auto predictions decreased to (at most) 5.2, 5.0, and 6.2%, respectively.

### D. Simulations

Simulations were obtained from an artificial listener (Sec. IIC and Appendix. A 4), using the same experimental set of noises from participants S01 to S12 and the same methods outlined earlier to derive ACIs. To distinguish the ACIs from the artificial listener from those of the participants, we refer to the first ones as ACISI. The ACISIs derived from the waveforms of participants S01-S03 are shown in Fig. 10. The remaining ACISIs, are shown in Fig. A.6.

**FIG. 10.**
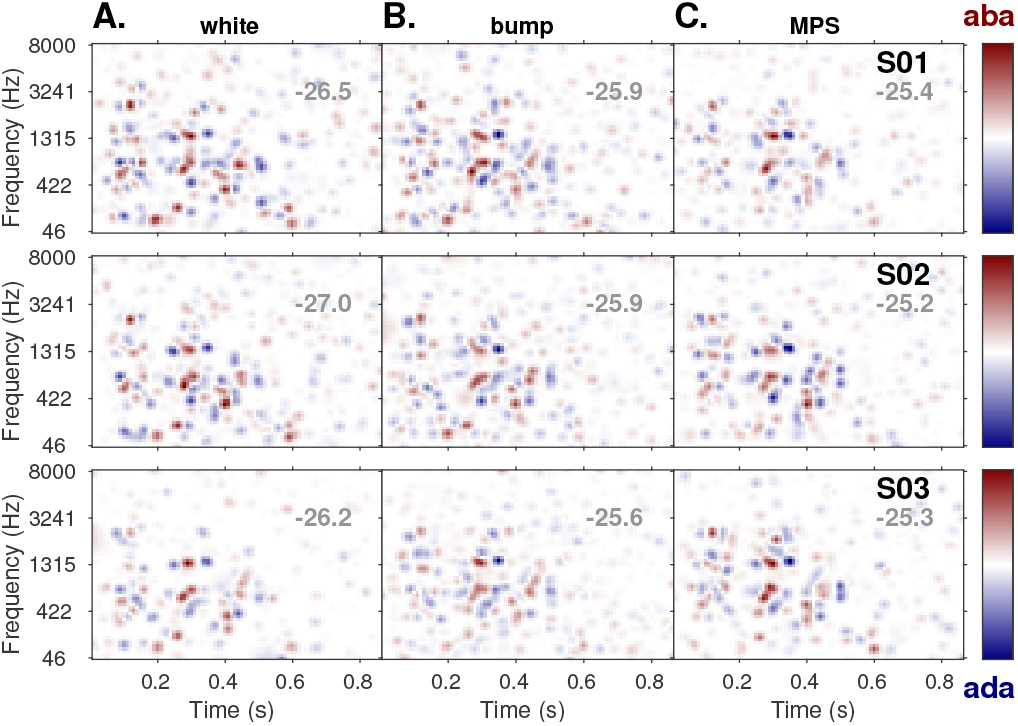
(Color online) ACIs derived from the simulations using the artificial listener (ACISIs) for (**A**) white, (**B**) bump, and (**C**) MPS noises using the set of noises from participants S01–S03 (top to bottom rows). The values in gray indicate the corresponding mean simulated SNR threshold expressed in dB. The ACIs derived from simulations for the remaining set of noises (S04–S12) are shown in Fig. A.6.

The ACISIs have more clusters of cues, compared to the experimental ACIs (Fig. 6). These clusters seem to be independent of the specific set of noises and are mainly located below 3000 Hz, and between *t* =0.1 and 0.5 s. All ACISIs have large weights in the *F*_1_ and *F*_2_ regions at *t* ≈0.3 s.

To quantify the similarity between ACISIs we assessed the cross predictions across “participants,” i.e., across noise sets of the same condition. These cross predictions, using ΔPA, are shown in Fig. 8**D**-**F**. The obtained ΔPA values were much higher than for the experimental ACIs (Panels **A**–**C**), ranging between 33.0 and 43.4% and were always significant. The significant cross predictions are enclosed by dashed pink boxes and are superimposed to all matrix elements in Fig. 8**D**–**F**.

The ΔPA cross predictions derived from exchanging ACISIs across noise conditions for the group are shown in Fig. 9**B**. Similar results were found within simulated individuals, as can be found in Fig. A.4, where all cross predictions were significantly above chance.

Based on the cross-prediction values averaged across data sets in Fig. 9**B**, the auto-predictions were 36.0, 37.6, 43.4% in the white-, bump-, and MPS-noise conditions, respectively. The high values of the cross predictions in the off diagonal, that differ by no more than 4.4% with respect to the on-diagonal values of Fig. 9**B**, again, support the similarity among the obtained ACISIs.

## IV. DISCUSSION

The objective of our study was to measure the token-specific effect of noise on the phonetic discrimination between the words /aba/ and /ada/. For this purpose, we predicted the listeners’ judgements using a microscopic (trial-by-trial) approach, the reverse-correlation method. The results allowed us to derive both macroscopic metrics of speech intelligibility and a microscopic characterization of the participants’ listening strategies by means of the time-frequency information in the ACIs.

We start this section by contextualizing the general performance results (Sec. IVA). We then focus on the interpretation of the microscopic ACI analysis to estimate the size of the token-specific effect of noise in our task, going through each of our study hypotheses (Sec. IV B– IV G). We conclude this section by indicating the limitations of the adopted approach (Sec. IVH).

### A. General performance in the task

The behavioral performance for each participant averaged across conditions was very similar, although there were participants with lower or higher overall performance, as indicated by the vertical shift of gray traces in Fig. 4 and as also seen in the SNR thresholds reported in Fig. 6, that ranged between −15.7 and −10.3 dB. When expressing the same data as a function of SNR we observed that, first of all and as expected, the difficulty of the task increased for lower SNRs. More precisely, the percent correct rates for /aba/ (Fig. 5**A**) and /ada/ trials (Fig. 5**B**) decreased from about 90% (for SNRs>−10 dB) to chance level (for SNRs< −17 dB). This strong effect of SNR was also visible using the discriminability index *d*′ (Fig. 5**C**) and the criterion c (Fig. 5**D**). The d’values were lower for bump and MPS noises than for white noises at any SNR. Given that all three noise types have approximately the same long-term spectrum (Fig. 2**B**) and thus should produce a similar EM effect, the lowered discriminability—that leads to an increased number of incorrect answers—can be attributed to the additional random fluctuations between 0 and 30 Hz in the bump and MPS noises (Fig. 2**D**, middle and right panels). In other words, the lowered discriminability can be partly linked to the MM effect and, as we discuss in the next subsection, to the increased token-specific effect in these two non-stationary maskers. Another observation that can be inferred from the lowered discriminability of bump and MPS noises is that there was a marginal, if any, effect of listening in the dips. In fact, a significant effect of this phenomenon should have resulted in better performance for more modulated maskers, an effect that we did not observe. In Fig. 5**D**, we observed a bias towards “aba” answers for SNRs below −15 dB, where the criterion values *c* were higher than 0, in contrast to the nearly constant c values for the SNR bins centered at or above −15 dB. In line with previous work on confusion patterns (Régnier and Allen, 2008), this might be an indication that below this level, some critical cue for discriminating /ada/ is no longer audible, while /aba/ is still correctly perceived. This is consistent with the argument that the acoustic cues for /d/ are typically located in a higher frequency range than that for /b/, being more likely to be effectively masked by white-noise-like maskers that have more energy in high frequency regions.

### B. ACIs and token-specific effect

The trial-by-trial (microscopic) analysis based on reverse correlation resulted in a total of thirty-six ACIs (Fig. 6), that characterized the listening strategy of the twelve participants in each background noise condition. In terms of prediction performance, the obtained ACIs were able to predict the categorical response in each trial (“aba” or “ada”) with a better-than-chance accuracy using 27 (out of 36) ACIs (Fig. 7**A**, open diamond). Additionally, the group results were significantly above chance (Fig. 7**A**, filled circles), indicating that the exact within-trial noise configuration had a significant influence on the participants’ responses or, in other words, that the three types of noises elicited a measurable token-specific effect. This effect was measured to be on average ΔPA=11% across all participants and conditions and ranged between 4.9 and 19.6% for the individual results (open diamonds in Fig. 7, diagonals in Fig. 8**A**–**C**). These results agreed with our preregistered expectations about the significance of ACI-based predictions and the size of their effect. Given that the GLM-fitted ACIs only relied on the T-F distribution of random noise envelope fluctuations, without using any explicit information from the targets, these results are supportive of Hypothesis H1 (Sec. I C).

The idea that different noise tokens produce a different amount of masking is not new. This has been shown in the context of, e.g., tone-in-noise detection (Ahumada and Lovell, 1971; Pfafflin, 1968), AM-in-noise detection (Varnet and Lorenzi, 2022), but also in psycholinguistic tasks (Varnet et al., 2013; Zaar and Dau, 2015). Using frozen noise, Zaar and Dau (2015) demonstrated that two particular white-noise tokens elicited different confusion patterns. They showed that, when presented with one specific frozen noise token at SNRs below 0 dB, the sound /gi/ was confused with /di/ or /bi/, but when the same sound was presented with a different frozen-noise token, /gi/ was robustly perceived for SNRs down to −15 dB.

A further exploration of the participants’ ACIs indicates that the random noise envelope fluctuations that can trigger a token-specific effect are concentrated in small but non-uniformly distributed T-F regions. These regions overlap with the position of acoustic cues in the target sounds, as emphasized in Fig. 6**G**-**I**. For example, the presence of a burst of energy in the random envelope fluctuations in the vicinity of the *F*_2_ onset will induce a response bias in favor of “aba” or “ada” depending on whether its spectral position matches the *F*_2_ onset frequency of /aba/ (1298 Hz) or /ada/ (1722 Hz). Similarly, subtle differences in the random envelope fluctuations in the region of the *F*_1_ onset, the *F*_2_ offset in the initial syllable, as well as in the plosive burst, located in the high-frequency region at the consonant onset, affected the listeners’ responses in a systematic way. The importance and relative weights of *F*_2_-transition cues and burst cues in the pereption of voiced plosive consonants has already been discussed in length elsewhere (e.g., Delattre *et al*., 1955; Ohde and Stevens, 1983) in particular in the presence of background noise (e.g., Alwan *et al*., 2011; Li *et al*., 2010). References to a possible role of *F*_1_ transitions are more seldom (Alwan *et al*., 2011; Delattre *et al*., 1955) but this cue was already found in our previous ACI studies (Varnet, 2015; Varnet *et al*., 2015a).

As noted above, there seems to be a correspondence between the T-F regions from Fig. 6, where the presence of random envelope fluctuations was particularly detrimental to the listener, and the acoustic cues from the targets. Arguably, this token-specific effect of noise can be seen as the counterpart of the MM effect described in Sec. I, as in both cases random envelope fluctuations induce confusions to the listeners, or a “sorting problem” using Drullman‘s words. For the MM effect, weak elements of the speech targets are confused with non-relevant noise envelope fluctuations. For the token-specific effect of noise, in contrast, it is the large random noise envelope fluctuations that are confused with relevant elements of the targets if the corresponding T-F locations overlap, affecting the listeners’ phonetic decisions. This is reminiscent of the conflicting cues that have been reported from the detailed analyses of phoneme confusions (Li and Allen, 2011; Régnier and Allen, 2008; Singh and Allen, 2012). These studies showed that some speech utterances present incidental acoustic cues that can be confused with characteristic cues of other phonemes, making the targets prone to confusions. In the case of our experiment, the conflicting cues are induced by the background noises.

### C. Token-specific effect in white noise

To estimate the token-specific effect of noise and dissociate it from the overall effect that is related to EM and MM, we used the metrics of out-of-sample prediction accuracy (Sec. II E 4 a) applied to the obtained ACIs. More specifically, we used the prediction accuracy (ΔPA) to measure the size of the effect and the cross-validated deviance per trial (ACVD_t_) to test its significance. Both metrics were compared to a null ACI, where the weights associated to the T-F noise distributions are set to zero, producing “aba”-”ada” predictions at the participant’s chance level.

The contribution of the token-specific effect to the overall effect in white noise was small but significantly above chance for 6 of the 12 participants (Fig. 7**A**, blue open markers), with an auto-prediction benefit ΔPA of 8.1% at the group level (Fig. 7**C**, blue filled marker). This estimate considers trials where conflicting cues misled the participants, as well as trials where the cues reinforced their correct answers. If we restrict this analysis to incorrect trials only, the auto-prediction benefit was significant for 7 participants (Fig. 7**B**, blue open markers) with an increased benefit of APAinc= 11.3% at the group level (Fig. 7**D**, blue filled marker). This small effect agreed with our preregistered expectations (Hypothesis H1), which were based on the few studies that have tried to disentangle between the different effects of noise (Drullman, 1995; Dubbelboer and Houtgast, 2007; Noordhoek and Drullman, 1997). These studies investigated the role of random envelope fluctuations, including MM and token-specific effects, using special types of signal manipulations to isolate EM from non-EM effects. Drullman (1995) and Noordhoek and Drullman (1997) performed listening experiments using vocoded sentences, and found that between 15% and 19% of errors can be attributed to the presence of random envelope fluctuations in the masker.Dubbelboer and Houtgast (2007) reached a somewhat similar conclusion for a word (consonant-vowel-consonant) recognition using steady-noise maskers. They were able to evaluate the effect of EM and non-EM independently using a wavelet-transform-based stimulus-resynthesis approach. Their results indicated that EM was the most detrimental type of interference, estimating a cumulative impact of all non-EM effects related to 19.7% of recognition errors.

Finally, a relatively small token-specific effect of noise or “noise-induced effect” was reported by Zaar and Dau (2015). The authors compared different sources of variability in the recognition of consonant-vowel-consonant words presented at different SNRs. In their analysis of source-induced variability they found that the variability in the background noises induced a significant perceptual effect, but that the effect was smaller than the variability in the speech sounds, between- and within-talkers. Although Zaar and Dau‘s analysis of within-participant variability was based on an analysis of confusion matrices that is not directly comparable to our out-of-sample metrics, their results are aligned with our ACI results.

### D. Token-specific effect in more fluctuating noises

The /aba/-/ada/ discrimination using “white-noise-like” bump and MPS noises with emphasized envelope fluctuations, was included to investigate a potential increase of the token-specific noise effect, as they were supposed to mask more efficiently the relevant elements of the speech targets. This is related to our preregistered Hypothesis H2 (Sec. I C). H2 is supported by the out-of-sample performance results, where ΔPA increased from 8.1% for white noises to 13.3 and 11.8% in the bump- and MPS-noise conditions, respectively (Fig. 7**C**, filled markers) and from ΔPA_inc_ = 11.3% to 18.5 and 17.0% for the corresponding analysis with incorrect trials only (Fig. 7**D**, filled markers). At the individual level, all (12 of 12) ACIs produced a significant prediction in the bump-noise condition and 9 of 12 ACIs did so in the MPS-noise condition (Fig. 7**A**, red and green open markers). In other words, with respect to the white noises, bump and MPS noises did not only lead to an increased MM effect as a consequence of their strong envelope fluctuations below 30 Hz (Fig. 2**D**), but they also led to an increased token-specific effect, with respect to the whitenoise condition.

### E. Between-subject variability

Due to the nature of our experimental task, that used two vowel-consonant-vowel words (/aba/ and /ada/) presented without semantic context, we expected that the ACI method should have resulted in globally similar listening strategies for all our participants. This was related to our preregistered Hypothesis H3 (Sec. IC). However, in contrast to this expectation, we obtained very heterogeneous ACIs (Fig. 6**A**-**F**). We assumed that two ACIs were globally similar when they could predict data significantly above chance when they were exchanged with each other. The results of this analysis were presented in Fig. 8**A**-**C**, where significant cross predictions using a specific ACI (shown along the abscissa) to predict the participants’ data (shown along the ordinate) are enclosed in dashed pink boxes. This analysis revealed that only 42, 67, and 57 (out of 132) cross predictions led to a performance that was significantly above chance in the white-, bump-, and MPS-noise conditions, respectively. This low number of significant cross predictions seems to be enough evidence to reject H3.

When analyzing the heterogeneity of the obtained ACIs (Fig. 6**A**-**F**) we did not find a direct link between the participants’ overall performance or any other information about them (e.g., language background, age, or audiometric thresholds) and the exact distribution of obtained T-F cues. For instance, the very good performing participants S04 and S08, who reached SNR thresholds below −15 dB showed significant cross predictions in all conditions, although the participants differ in their linguistic background. In another example, the bump ACIs from S03 and S10 produced significant cross predictions despite the difference in overall SNR threshold of 3.2 dB during the experiments, while we did not find a significant similarity in their strategy for the other two noise types.

Differences in listening strategies were also reported by Singh and Allen (2012), who observed a non-negligible between-subject variability in the perception of noisy /b/ and /d/ sounds compared to, e.g., /t/ and /g/. They related this finding to the observation that /b/ and /d/ involve multiple cues (Alwan et al., 2011; Dorman et al.,1977) unlike /t/ and /g/ which have an identifiable single feature that makes them noise robust (Li et al., 2010). Similarly, for our task, the redundancy of available cues noted in Sec. IVB may have enabled our participants to use different listening strategies, as supported by visual inspection of the obtained individual ACIs (Fig. 6). The most logical explanation for such “disparity” in the use of cues may be due to the diversity of listeners’ linguistic backgrounds (e.g., Pallier et al., 1997), however, this contrasts with results of other studies where cue disparities have also been found in participants with a common linguistic background (Clayards, 2018; Singh and Allen,2012; Zaar and Dau, 2015). Our results do not provide any evidence supporting any of these two possibilities. Further studies are required to investigate the origin of the inter-individual variability observed in the ACIs.

### F. Artificial listener

In this study, the artificial listener was used as a baseline for human performance, under the assumption that measurable changes in auditory-model responses due to changes in the signals—for us, when using different noise maskers—reflect an effect that might be observable by human listeners (Green and Swets, 1966). More concretely, the artificial listener was used to confirm that (1) a higher out-of-sample prediction is reached for more fluctuating noises (bump and MPS noises) with respect to the steady-state white noises, and (2) the specific set of generated noises of the same type does not influence significantly the obtained ACIs. This last point is important because the algorithms for the noise generation were developed and adjusted to particularly influence the modulation frequency content below about 30 Hz and we wanted to confirm that these manipulations do not bias a specific set of responses in the “objective” auditory model decision. At the same time, we expected that the decisions of the artificial listener should elicit a measurable token-specific effect due to the trial-by-trial (microscopic) nature of the template-matching approach.

The results of the simulations were presented in the bottom panels of Figs. 8–9 (out-of-sample metrics), Fig. 10 and Fig. A.6 (obtained ACISIs). All the obtained ACISIs produced a measurable token-specific effect of noise with predictions significantly above chance. The average token-specific effect using ΔPA, averaged across the 12 data sets, was estimated to be 36.0, 37.6, and 43.4% for white, bump, and MPS noises, respectively. These results confirm that a reliable token-specific effect can be measured using the artificial listener and that a mild but systematic increase in ΔPA was observed for the bump and MPS noises with respect to the white noises, supporting the validity of Hypothesis H4 (Sec. IC). Furthermore, the significance analysis showed that all auto and cross predictions produced significant out-of-sample metrics (see the pink dashed squares in Fig. 8**D**-**F**), supporting the statement—also contained in H4—that the specific set of noises did not influence the ACI_sim_ estimation. Despite this support to H4, we observed that the ACISIs contain many more T-F cues than the experimental ACIs, suggesting that the underlying strategies between “this” artificial listener and the participants are differently weighted and that, on average, less T-F cues were used by the participants.

### G. Token-specific effect as a form of informational masking

As discussed in Sec. I, the token-specific effect is not encompassed in the traditional definition of EM and MM effects. In this section we argue that the token-specific effect is a form of informational masking (IM).

IM has been defined using different, partly-overlapping notions proposed over decades of research. While IM effects are often defined as any masking effect that cannot be attributed to EM, or as a masking occurring at a “central” level, these simple definitions have been criticized for their lack of precision (e.g., Durlach et al., 2003; Watson, 2005). In particular, from these perspectives, it is unclear whether MM should be identified as EM or IM (Conroy and Kidd, 2021; Durlach et al.,2003). For these reasons, several authors have proposed alternative and more operational definitions of IM that have led to a trichotomy between EM, MM, and IM effects (Durlach, 2006; Stone et al., 2012). According to these more refined views, IM effects are responsible for a reduction in perceptual thresholds due to (1) masker uncertainty, or (2) similarity between masker and target.

IM is often linked to the notion of masker uncertainty (Alexander and Lutfi, 2004; Micheyl et al., 2000; Neff and Green, 1987), with higher levels of uncertainty maskers causing larger IM effects. In tasks as the one presented in this study, uncertainty arises from the trial-by-trial variability induced by random envelope fluctuations in the noise maskers. Contrary to EM and MM effects, which reflect the inaudibility of some critical cues from the speech target due to the “saturation” of the corresponding cochlear- or modulation-frequency channels, the token-specific effect results from the sensitivity of listeners to fine T-F details in the masker waveforms, that are susceptible to be confused with relevant speech cues. In this respect, our /aba/-/ada/ experiment can be regarded as a detection of specific acoustic cues in the presence of randomly-occurring conflicting cues, a task that is reminiscent of classic IM experimental paradigm with uncertainty introduced in the spectral (Alexander and Lutfi, 2004), temporal (Conroy and Kidd, 2021), or spectro-temporal domains (Kidd et al., 2002). This parallel may seem surprising as researchers studying IM effects have often considered broadband stationary noise (as our white-noise masker) as a low-uncertainty condition (Lutfi et al., 2003), using it as a no-IM baseline (e.g., Agus et al., 2009; Neff and Green, 1987). However, other authors have considered the possibility that a relatively small amount of uncertainty can already induce some form of IM (Goossens et al., 2008; Lutfi, 1990), which is in line with our findings.

Previous work has indicated that masker-target similarity influences the amount of IM masking. This can be evidenced, for instance, using maskers that share some lexical or phonetic information with the speech targets, as in speech-in-speech tasks (Brungart, 2001; Hoen et al.,2007). This is obviously not the case for the three types of maskers investigated in our study. However, given that the derived ACIs were obtained from the information contained in the noises only, it can be inferred that random envelope fluctuations from these maskers induce small but consistent perceptual biases towards one or the other response alternatives. In turn, this indicates that the conflicting cues present in the noise did not just provide a distraction, but were actually confused with relevant speech cues from the target. In this sense, we can consider that the maskers *do* share information with the speech target at the acoustic cue level: when the noise token presents some acoustic characteristics similar to one of the two targets, by chance, it is more likely to elicit the corresponding response.^5^

More recently, uncertainty and similarity have been merged into the notion that IM effects originate from a more general source-segregation problem (Kidd et al.,2002; Lutfi et al., 2013; Shinn-Cunningham, 2008). Furthermore, several studies have revealed large individual differences in susceptibility to IM, that may reflect differences in listening strategies (Alexander and Lutfi, 2004; Durlach et al., 2005). In line with these results, as highlighted in Sec. IVB, the interference of noise envelope with the perception of a speech target can be seen as a “sorting problem,” because listeners erroneously use the confounding cues from the masker as part of the speech target. These ideas seem to offer an explanation to the large non-negligible variability in our estimates of the token-specific effect, that led us to reject Hypothesis H3. In other words, the measured token-specific effect using the ACI approach is reminiscent of classic IM definitions, as the ACIs suggest the existence of a processing that is modulated by a central (top-down) prior that may differ from participant to participant.

### H. Limitations of the approach

The prediction performance of our ACI approach was used to quantify the token-specific effect of noise on an /aba/-/ada/ discrimination. For this quantification, we assumed that the participants’ responses could be predicted purely based on the random envelope fluctuations of the noises that were used to mask the target words. Under this strict assumption, we believe that the reported performance metrics represent only a lower bound of the actual token-specific effect because:

1. We considered the envelope of noise-alone waveforms instead of the envelope of the noisy speech sounds. We assumed this to ensure that the estimated token-specific effect came from the noise maskers and not from the speech targets themselves. Nevertheless, given that envelope extraction is a highly nonlinear process, this assumption implies that the effect of spurious modulations arising from speech-noise interactions (Dubbelboer and Houtgast, 2008; Stone *et al*., 2011) is negligible.
2. The transformation of noise waveforms into T-F representations—the inputs to the GLM—is based on a set of linear cochlear filters followed by a simplified envelope extraction (e.g., Osses *et al*., 2022), ignoring the potential influence of more central stages of auditory processing on the estimated token-specific effect. In this study, we decided to keep the T-F transformation as simple as possible.
3. Following a similar principle of simplicity as in the T-F transformation, the GLM approach we used as a statistical model back-end to relate noise (T-F) representations with the participants’ responses did not consider interactions between GLM predictors. It is possible, however, that listeners make their decisions based on a non-linear combination of cues.

## V. CONCLUSIONS

In this study, we conducted a microscopic (ACI) analysis of participants’ responses in an /aba/-/ada/ discrimination task, using three different white-noise-like and contextless maskers. We demonstrated that:

1. The detailed noise structure has a measurable effect on a phoneme-in-noise discrimination task. A particular noise token can bias the participants’ choice towards one alternative or the other depending on its exact time-frequency (T-F) content. We argued that this token-specific effect of noise is a form of informational masking (IM) that can be elicited by any random masker, including white noise.
2. At low SNRs (≈ −14 dB), this effect accounts for at least 8.1% of the participants’ responses in white noise (or 11.3% of the errors). When considering other maskers that have larger amounts of random envelope fluctuations, this percentage increased to 13.3% (or 18.5% of errors) and 11.8% (or 17.1% of errors) for the bump and MPS noises, respectively.
3. Substantially similar results were obtained using an auditory model that is based on a microscopic (template-matching) approach. The model was used to simulate the same /aba/-/ada/ discrimination task as our study participants. In this case, the token-specific effect of noise was estimated to be between 33.0% and 43.4% of the the model correct responses. Based on these results, the model adopted, as expected, a more optimal and consistent decision strategy, given that the model relied on more T-F cues than our participants with better (and always significant) cross predictions.
4. Contrary to Hypothesis H3 (Sec. IC), we observed a large variability in listening strategies, both between participants and between masker types. A close investigation of the results revealed that, although the primary F2 cue is seen in almost every individual ACI, the weights attributed to secondary acoustic cues appear to differ between participants.

## ACKNOWLEDGMENTS

We would like to thank Richard McWalter for his critical reading of earlier versions of this study and Christian Lorenzi and the rest of the “Modulation Group” for the many discussions during the course of this project. This study was supported by the French National Research agency through the ANR grants “fastACI” (Grant No. ANR-20-CE28-0004) and “FrontCog” (Grant No. ANR-17-EURE-0017).

## FOOTNOTES

1 Noise maskers are known to degrade target sounds in terms of temporal fine structure (e.g., Drullman, 1995) and temporal envelope. In the present paper we focus on degradations due to temporal envelope information, which are typically assumed to have a larger impact on speech perception (e.g., Shannon *et al*.,1995).

2 The DC removal was applied to the absolute value of the Hilbert envelope, just before applying the FFT. The DC removal was applied to better visualize the spectrum levels in Fig. 2**C**. Without this processing, the DC amplitudes were going to have an amplitude of approximately 65 dB for all three noise types.

3 We tested different sets of Gaussian-pyramid parameters, exploring the required number of levels and the inclusion of level 0 (i.e., the inclusion of the original 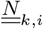 matrix). The specific configuration of the pyramid did not seem to affect critically the overall shape of the resulting ACIs.

4 To get an indication of the level of ΔPA that can be attributed to chance only, we assumed that the predictions in each set of 4000 noises follow a binomial distribution ~ *B*(4000, 0.5), i.e., we assumed that the success of the prediction is determined by chance with *P*(*r_i_* = “aba”) = 0.5. Considering the one-sided 95% confidence interval, PA needs to be equal to or greater than 51.3% (or ΔPA≥ 2.6%, after correcting for guessing). This boundary is increased to 52.39% (ΔPA≥ 4.78%) for the analysis of incorrect trials, where only 29.3% of the trials are used (B(1172, 0.5)). This “significance test” should only be considered as referential because (1) due to data exclusion, the number of trials is reduced by ≈10%, so 4000 and 1172 are not the exact numbers that should be used in the binomial approximation, and (2) the probability of successful prediction by chance deviates slightly from 0.5 depending on the exact ratio of “aba” and “ada” responses in the participant data. To avoid unnecessary confounds, we refrained from including the exact number of trials and chance levels, and we just presented the estimated chance boundary as a visual aid in Fig. 7.

5 In line with this interpretation, several participants reported being very confident in their answer to some trials, and very surprised to find out that their response was in fact incorrect. This is the kind of perceptual experience that we would expect to see if the masker was indeed confused with the speech targets.

## APPENDIX A SUPPLEMENTARY MATERIALS

### 1. Participants’ details

Twelve participants took part in our study aged between 22 and 43 years old. Further details of the participants are given in Table I. We characterized their hearing status by measuring audiometric thresholds and their performance in a speech-in-noise test, whose details are given next. While the hearing thresholds were used as the only inclusion criterion, the speech-in-noise thresholds were planned to give an indication of the participants’ supra-threshold hearing status, to be used as referential data for the design of future studies.

**TABLE I.** Participants’ details. The age is expressed in years at the time of testing. Participants S06 and S12 were the two last participants to complete the experimental sessions. Their data were excluded in Sec. A 3 b for the analysis with the preregistered number of *N* = 10.

| Subject | Age | Gender | Mother tongue | Speaks French |
| --- | --- | --- | --- | --- |
| S01 | 33 | M | French | Yes |
| S02 | 36 | M | Spanish | No |
| S03 | 31 | F | French | Yes |
| S04 | 38 | M | French | Yes |
| S05 | 24 | F | Italian | No |
| S06 | 43 | M | French | Yes |
| S07 | 23 | M | French | Yes |
| S08 | 27 | F | Turkish | Yes |
| S09 | 25 | M | French | Yes |
| S10 | 22 | F | French | Yes |
| S11 | 36 | M | Spanish | No |
| S12 | 22 | M | French | Yes |

#### a. Audiometric thresholds

Audibility thresholds were measured using pure-tone audiometry at six frequencies (250, 500, 1000, 2000, 4000, and 8000 Hz) and had average thresholds between 0.8 (S07) and 12.5 dB HL (S02) in their best ear, meeting our inclusion criterion of having thresholds of 20 dB HL or better. The obtained hearing thresholds are shown in Fig. A.1.

#### b. Intellitest

The participants’ supra-threshold hearing status was measured using the Intellitest speech-in-noise test (Gnansia *et al*., 2014). The Intellitest is a closed-set speech material of 16 words of the structure VCVCV containing three takes of each word (total of 48 samples). The dataset was split into three single lists of 16 non-repeated words. Three single lists were evaluated twice, either using a speech-shaped noise (SSN), or using an 8-Hz amplitude modulated version of the SSN. In these experiments the speech level was adjusted targeting a 50% score. The threshold estimate for each noise condition obtained from the median of the three single list runs in each condition are shown in Fig. A.2. In this figure we grouped the participants into native French speakers (*N* = 8, blue traces, “French”) and the rest of the participants (*N* = 4, red traces, “Non-French”). The speech reception thresholds (SRTs) for the French group had median thresholds of −10.4 and −27.7 dB in the steady-noise and 8-Hz AM noise, respectively. The threshold using the modulated masker was 17.3 dB lower (better) than the threshold in the steady-noise condition. The results for non-French group were −7.3 and −23.1 dB in the steady-noise and 8-Hz AM noise, respectively. These thresholds were higher (worse) than the thresholds obtained for the French speakers, by 3.1 and 4.6 dB for the two noise conditions. The difference between performance in steady and modulated noise was 15.8 dB.

**FIG. A.1.**
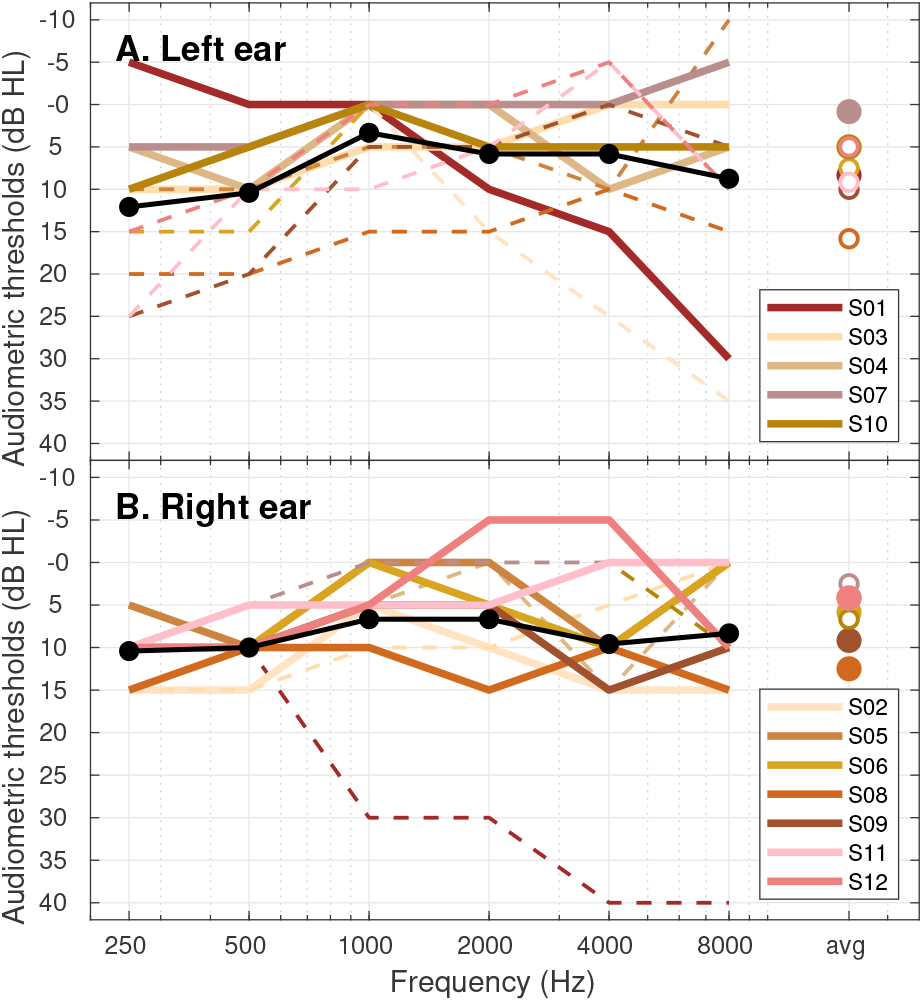
Audiograms for all participants. Left and right ear thresholds are shown in Panels **A** and **B**, respectively. The participant’s best-ear thresholds are connected by continuous traces, and the subject ID is indicated in the corresponding panel legend. Average thresholds across participants are indicated by the black traces and the average audiometric threshold for all tested frequencies between 250 and 8000 Hz are indicated by the right-most markers (filled symbols are used when those averages are from the participant’s best ear).

**FIG. A.2.**
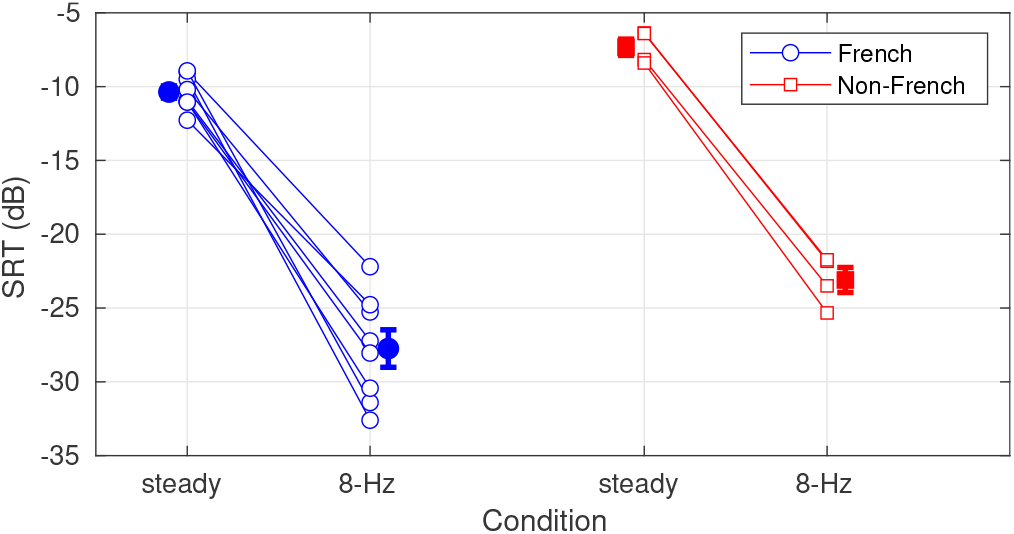
Results for the evaluation of the Intellitest speech material using a steady SSN background noise or an 8-Hz 100% amplitude-modulated version of it. We present separately the results for French speakers (blue) and non-French speakers (red). The filled symbols indicate the group mean thresholds and the error bars represent one SEM.

### 2. Retrieving the experimental sounds stimuli

The thirty-six sets of noises and the two speech samples (/aba/ and /ada/) used in the experiments can be retrieved either from Zenodo (Osses and Varnet, 2022c) or using our in-house fastACI toolbox. To retrieve the sounds using the toolbox, the script `publ_osses2022b_-preregistration_0_init_participants.m` needs to be run. Note that for recreating the MPS noises, the PhaseRet toolbox (Průša, 2017) needs to be installed and compiled. No extra dependencies are required to reproduce the white and bump noises.

Once generated, the noises will be stored in separate folders named NoiseStim-white, NoiseStim-bump, and NoiseStim-MPS, each of them containing 4000 waveforms using a numbered labeling (`Noise_0000ĩ.wav-Noise_-04000.wav`).

### 3. Complementary information to the data analysis

#### a. Analysis using the data of all participants: Extra figures

Figure A.3 is complementary to Fig. 8 and contains the cross-prediction values using the deviance per trial (CVDt). This information had been omitted in Fig. 8. The panels (**A**–**C**) show the CVD_t_ values obtained from the experimental data, whereas the bottom panels (**D**–**F**) show the corresponding values for the artificial listener. In this case, the insets ‘S01’ to ‘S02’ indicate the set of waveforms used to run the fixed normal-hearing auditory model (Sec. A4).

Figure A.4 is complementary to Fig. 9 and shows the individual 3-by-3 matrices for each participant for the cross predictions between noises. The top and bottom panels show ΔPA values derived from the experimental data and from the artificial listener, respectively. The arithmetic average of ΔPA values across participants, corresponds to the 3-by-3 matrix presented in Fig. 9.

Figure A.5 (top panels, **A**–**C**) shows the correlations across ACIs (from Fig. 6). Figure A.5 (middle panels, **D**–**F**) shows the correlations across ACISIs (from Fig. 10 and A.6). The global results in both rows of panels is similar to the results obtained using the ΔPA metric: The correlations across experimental ACIs (top panels) are lower than the correlations across ACISIs obtained from the simulations. The off-diagonal correlations are 0.33, 0.20, 0.29 for white, bump, and MPS noises in the top panels. The corresponding values in the middle panels are 0.70, 0.70, and 0.74. The results in the bottom panels indicate the average results for the Pearson correlations within participant but between noises, comparable to Fig. 9. In agreement with Fig. 9, the off-diagonal correlations had an average of 0.33 (Panel **G**, experimental data) and 0.69 (Panel **H**, simulation data).

**FIG. A.3.**
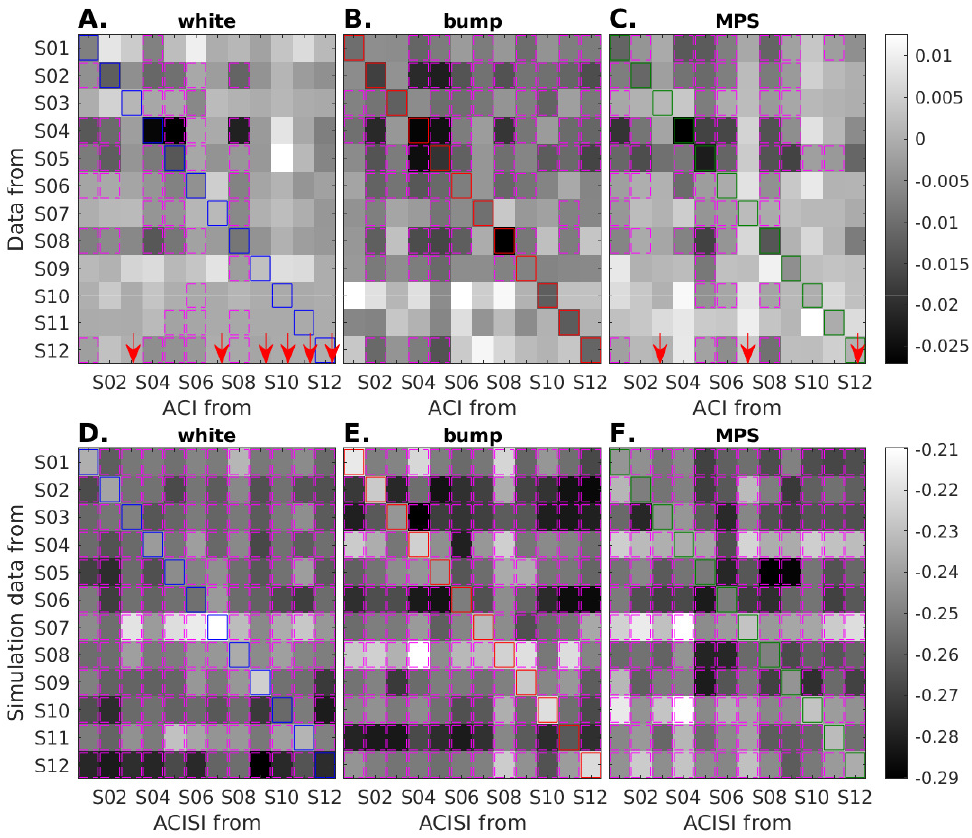
(Color online) **A**–**C**: Between-subject cross-prediction matrices for the three conditions using CVD_t_. The matrices contain the deviance benefit plus 1.64 SEM (ΔCVD_t_+ 1.64 SEM). When this quantity is less than 0, the cross predictions using the ACIs from the participants indicated in the abscissa are able to predict significantly above chance the data from the participants indicated in the ordinate. Those cases are enclosed in pink boxes. The main diagonals are enclosed in colored squares and correspond to the same auto-prediction values that are shown as open markers in Fig. 7**A**. The red arrows indicate the ACIs that did not achieve significant auto predictions. In such a case, the significance of the cross predictions was not evaluated. **D**–**F**: Same as the top panels, but using the ACISIs derived from the artificial listener.

#### b. Analysis using the data of ten participants

Here we replicated the reported mixed ANOVAs and the assessment of group averaged performance excluding the data of the two last completed participants (S06 and S12), i.e., only using data from the preregistered number of participants *(N* = 10).

##### **Behavioral performance** (as in Sec. IIIA)

###### Two-way mixed ANOVA on SNR

This analysis was run to test the learning effect on SNR (comparable results as with N = 12). There was a significant effect of the factors masker (*F*(2,287) = 15.82, *p* < 0.001) and test block (*F*(1,287) = 33.06, *p* < 0.001). As with *N* = 12, a post-hoc analysis confirmed that the effect of masker type was due to a difference in the bump-noise condition compared to the other two types of noise, with white and MPS noises having statistically the same SNRs.

**FIG. A.4.**
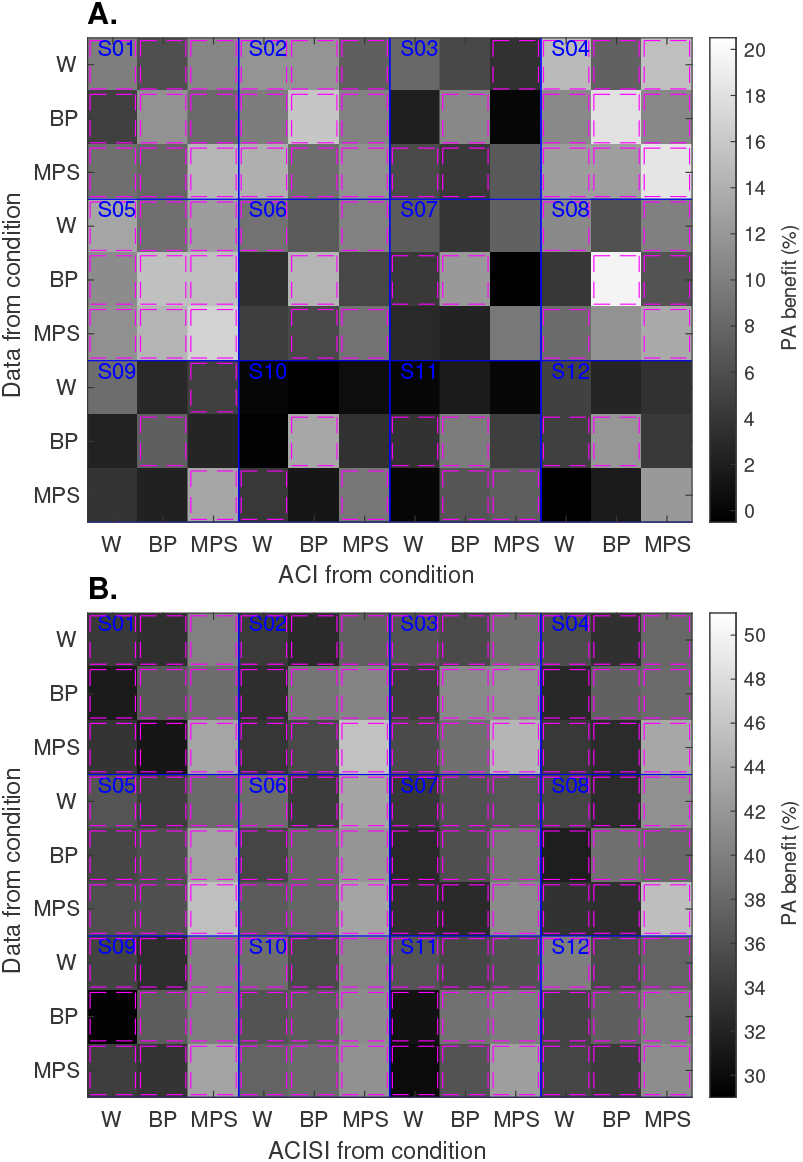
(Color online) Between-noise cross-prediction (3-by-3) matrices for (**A**) each participant and for (**B**) each artificial listener, using ΔPA. The pink boxes indicate cross predictions that provided significant better-than-chance ΔPA values. In each 3-by-3 matrix, the main diagonal takes values that were overall higher than those from the off-diagonals.

**Two-way mixed ANOVA on** *d*′ (comparable results as with *N* = 12): There was a significant effect of the factors masker (*F*(2,137) = 10.24, *p* < 0.001) and SNR (*F*(1,137) = 788.09, *p* < 0.001).
**Two-way mixed ANOVA on** *c* (comparable results as with *N* = 12): There was a significant effect for the factor SNR (*F*(1,137) = 13.12, *p* < 0.001), but not for the factor masker (*F*(2,137) = 1.41, *p* = 0.249).

###### **Out-of-sample prediction accuracy** (as in Sec. IIIC)

This analysis is comparable to the results shown in Fig. 7. The group results for the ΔPA metric are 8.4, 13.3, and 12.0%, for the white, bump, and MPS noises, respectively. For the analysis of incorrect trials only, the corresponding values were 12.3, 18.3, and 16.9%.

#### 4. The artificial listener

As briefly described in Sec. IIC, an auditory model was used to simulate the performance of an average normal-hearing listener who uses a fixed decision criterion to compare sounds. In this sense, the model is used as an artificial listener.

**FIG. A.5.**
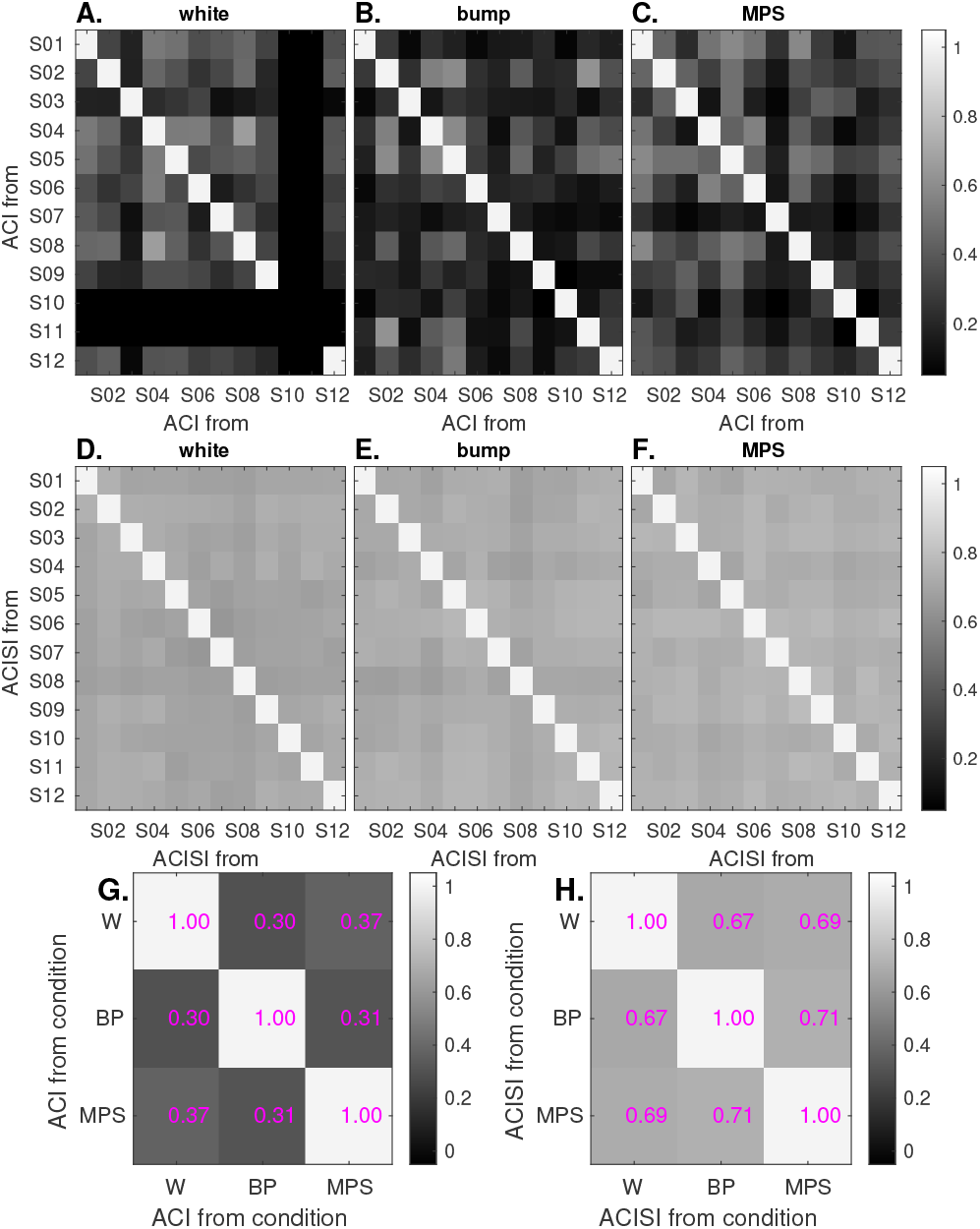
Pearson correlation values for ACIs between participants obtained from the experimental data (top panels, **A**–**C**) or obtained from simulations (middle panels, **D**–**F**). The correlation values in the bottom row were obtained from the ACIs between noise conditions for each participant, and then the values were averaged across participants. All matrices in this figure are symmetric with respect to their diagonal.

##### a. Model description

The model consists of a front-end and a back-end processing. The front-end processing converts an incoming sound waveform into an internal representation, i.e., into a representation that is believed to reflect how sounds are actually transformed along the ascending auditory pathway (e.g., Osses *et al*., 2022).

###### a. Front-end processing

The auditory model accepts monaural input waveforms and delivers a three-dimensional signal in time, audio frequency, and modulation frequency, that are expressed in model units (MU), an arbitrary amplitude unit (e.g., Kohlrausch *et al*., 1992). Most of the model stages have been previously described in detail (Osses, 2018;Osses and Kohlrausch, 2018, 2021). Here, we provide a short description of each stage, emphasizing some small implementation updates.

###### Outer- and middle-ear filtering (updated)

Two cascaded 512-tap FIR filters are used to produce a combined bandpass frequency response (Osses *et al*., 2022, their Fig. 3). In contrast to the previous model version (Osses and Kohlrausch, 2021), the middle-ear filter is implemented using the linear-phase version instead of its minimum-phase implementation. A group delay compensation is applied to the filtered signal.

**FIG. A.6.**
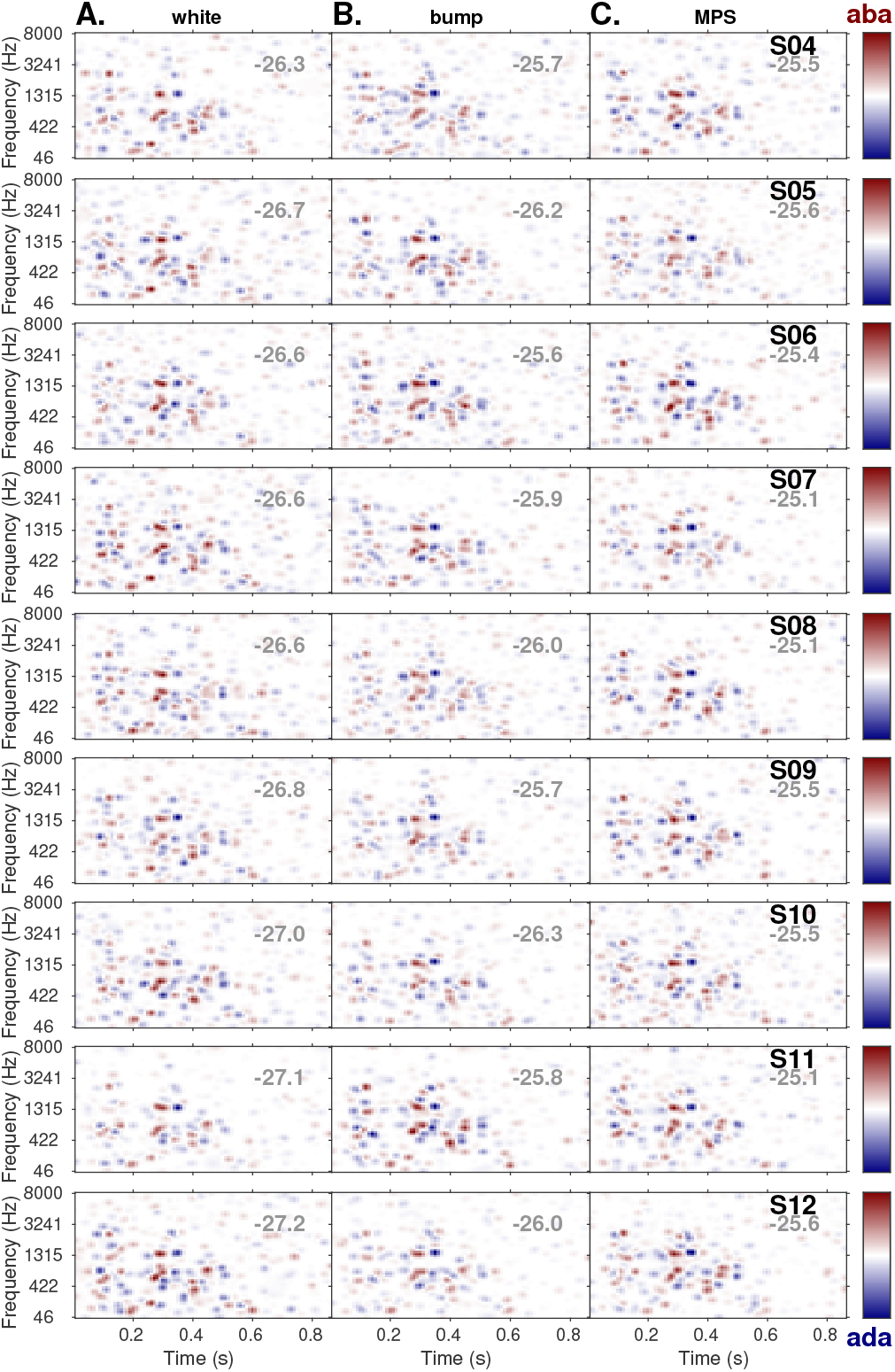
ACIs derived from the simulations using the artificial listener (ACISIs) for white (column **A**), bump (column **B**), and MPS noises (column **C**) using the set of noises from participants S04–S12 (top to bottom rows), which were not shown in Fig. 10. The values in gray indicate the corresponding mean simulated SNR threshold expressed in dB.

###### Gammatone filter bank

Set of 31 audio frequency bands with *f_c_* between 86.9 Hz and 7819 Hz, spaced at 1 ERB_*N*_, as described by Hohmann (2002). Only the real part of the complex-valued outputs of the filter bank is used.

###### Half-wave rectification and LPF

The half-wave rectification is followed by a chain of five cascaded first-order IIR filters with *f*_cut-off_= 2000 Hz. This chain produces a filter response with a —3-dB point at 770 Hz.

###### Adaptation loops

This stage approximates the effect of auditory adaptation at the level of the auditory nerve by using five feedback loops based on a resistor-capacitance analogy (full details in Osses and Kohlrausch, 2021, App. B) with time constants *τ* = 5, 50, 129, 253, and 500 ms. We used the parameter configuration indicated by Osses and Kohlrausch (2021) that uses a limiter factor of 5 instead of 10.

**FIG. A.7.**
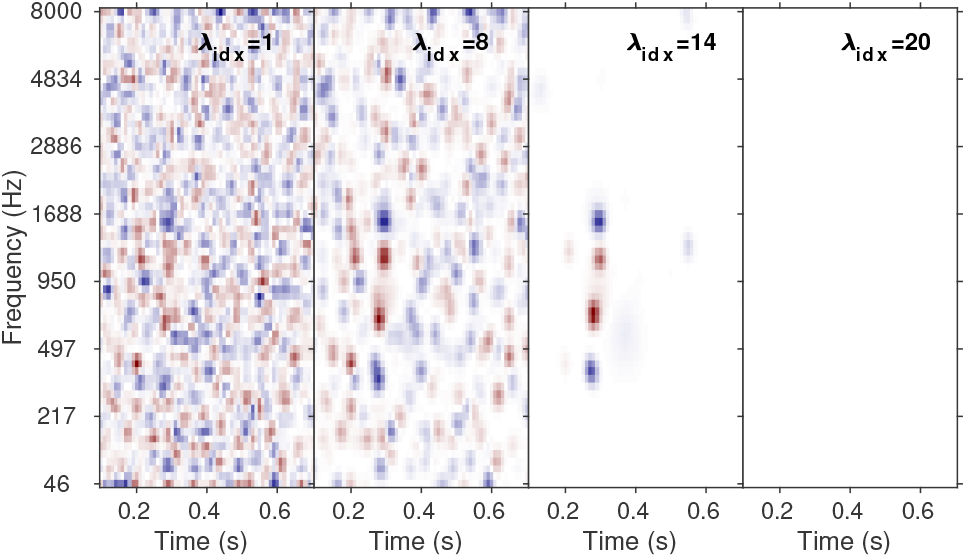
ACI for participant S01 for the MPS condition using different hyperparameter λ values. During the fitting procedure, the higher the λ value the fitting procedure looks for less and smoother time-frequency cue candidates. The right-most ACI corresponds to the null ACI, where the only non-zero parameter is the intercept *c_k_*. The lambda values in this figure range between λ_1_ = 1.1 · 10^-3^ and λ_20_ = 0.1.

###### Modulation filter bank (updated)

The implementation was mainly based on the filter banks by Osses and Kohlrausch (2021) and Jepsen et al. (2008). However, (1) the first-order 150-Hz LPF was implemented as an attenuation gain (see, Osses and Kohlrausch, 2021, their Fig. 14C), and (2) the filters were designed using a Q factor of 1, resulting in 7 modulation filters centered at 2.5, 5, 10, 25, 75, 225, and 675 Hz.

###### b. Back-end stage

The auditory model performed the same experimental paradigm as each of the twelve participants for the three noise conditions. For the simulations, the same order of noise presentation and the same level roving as the participants was used, but the exact SNR in each interval depended on the specific model responses.

To generate a decision outcome the internal representation of the current trial *R_c_*—the output of the frontend processing—was compared with the /aba/ (*T*_1_) and /ada/ (*T*_2_) template, derived at a supra-threshold SNR of −6 dB (i.e., with the speech sample presented at a level of 59 dB SPL). The comparison was based on a cross correlation at lag 0. The artificial listener indicated the option “aba” if *R_c_* · *T*_1_ ≥ *R_c_* · *T*_2_ + *K* or the option “ada” if *R_c_* · *T*_1_ < *R_c_* · *T*_2_ + *K* (Osses and Varnet, 2021). More formally:

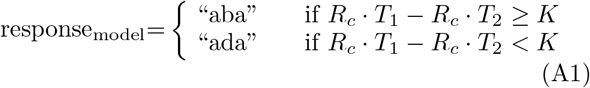

###### b. Calibration of the model

The bias *K* depended on the exact set of templates and on the type of noise. The use of a *K* = 0 led to biased model responses. For this reason we used *K* as a free parameter. The fitting of this parameter was performed before the simulation of each new set of noises, using a constant stimulus procedure at a very low speech level arbitrarily set to an SNR of −40 dB (i.e., at a speech level of 25 dB SPL)—a condition that the model should not be able to solve—and we stored the cross-correlation values (CCV = *R_c_* · *T*_1_ – *R_c_* · *T*_2_) for all 4000 trials. The final *K* value was chosen to be the median of the CCVs.

###### c. ACIs derived from simulations

The ACIs derived from simulations that used the set of noises of participants S01–S03 were shown in Fig. 10, in the main text. The remaining ACIs that used the set of noises of participants S04–S12 are shown in Fig. A.6.

#### 5. ACIs for different hyper parameter values

The time-frequency weights in the ACI_*k*_ and intercept *c_k_* are obtained during the GLM fitting procedure (see Sec. IIE 3), using the noise vector *<u>N</u>_k,i_* and the participant’s (or artificial listener’s) responses. During the 10-fold cross-validation procedure of the lasso regression, different hyperparameter values are evaluated. The intermediate ACIs obtained for four different values of the hyperparameter λ applied to the data of participant S01 in the MPS condition are shown in Fig. A.7. The rightmost ACI, the null ACI, is particularly important for the prediction performance that we used, because the goodness-of-fit metrics of CVD_t_ and PA (see Sec. IIE 4 a) were referenced to that null ACI, whose performance was nearly close to chance. An additional scaling was applied to the PA metric, to correct for guessing, with expected ΔPA values between 0% (performance at chance according to the null ACI) and 100%.

